# The Cooperation Ladder: Scale-dependent payoffs and population dynamics create surges, stalls and reversals

**DOI:** 10.1101/2021.02.19.432029

**Authors:** Eric Schnell, Robin Schimmelpfennig, Michael Muthukrishna

## Abstract

Human societies have expanded from small bands to large nation-states over the past 12,000 years. Yet, how groups scale up cooperation and why cooperation varies widely between societies, remains a central puzzle. We present a theoretical model that addresses these puzzles by extending the classic Stag Hunt game to (a) multiple players, (b) multiple rewards (“stags”) of different sizes, and (c) endogenous population growth. This framework reveals a “cooperation ladder” where each rung corresponds to a reward that requires a threshold number of cooperators. As cooperation increases, larger rewards become attainable. Securing a larger reward raises carrying capacity (e.g. by providing more food or energy), enabling subsequent population growth and unlocking the possibility of further cooperative gains. However, between these thresholds, cooperation can stagnate or reverse, effectively incentivizing free-riders at intermediate levels. We show that history matters. Early cooperation and population growth can set groups on divergent, path-dependent trajectories. Our model predicts multiple stable equilibria, with surges in cooperation when a new threshold is within reach, and stalls when higher rewards seem unattainable. This framework helps explain key patterns including why cooperation can sometimes accelerate rapidly, why some societies get stuck at smaller scales, and how seemingly selfish behavior can persist in cooperative groups. Expanding the scale of cooperation may depend on how incentives correspond to both environmental conditions, existing levels of cooperation, and population dynamics, offering a new lens on historical transitions in social complexity and insights for modern coordination challenges such as climate change.

**Significance Statement:** Small groups can sometimes grow into large-scale cooperative societies, while other societies stagnate at smaller scales. Our theoretical model reveals that population growth and cooperative thresholds for accessing resources or energy create a “cooperation ladder.” As a group’s population increases, new larger-scale payoffs become attainable, which in turn fuels further growth. However, between these critical thresholds, cooperation tends to stall and some free-riding is tolerated. This mechanism offers a new explanation for historical bursts in population and cooperation (such as the agricultural and Industrial revolutions), why some societies remain small, and how selfish behavior can persist in cooperative groups. It also provides insights into global coordination challenges like climate change by highlighting the material conditions needed to sustain and expand large-scale cooperation.

## Introduction

The evolution of human cooperation from small bands of related individuals to large societies of anonymous strangers remains one of science’s biggest puzzles. Two decades ago, *Science* identified the question “How did cooperative behavior evolve?” among its top 25 unsolved problems (1). Significant progress has been made in identifying mechanisms that sustain cooperation, from the role of kinship and inclusive fitness to reciprocity, norms and reputation, religion, and institutions (2–6). Yet understanding these mechanisms raises a deeper question: How do societies transition from small-scale to large-scale cooperation, given that different scales often rely on entirely different mechanisms? In other words, what drives the expansion from villages where people are related (kin selection) and everyone knows each other (reciprocity and reputation) to a nation or alliance of millions cooperating via impersonal institutions? Understanding how large-scale cooperation emerges is critical for tackling 21st-century challenges (e.g. climate change, pandemics) that demand global coordination.

Historical evidence shows that the scale of cooperation has increased dramatically over millennia, particularly since the rise of agriculture, Industrial Revolution, and Green revolutions (7–9). Over this timescale, rates of violent conflict have generally declined and large supra-national cooperative structures from expansive trade networks to NATO and the United Nations have emerged (10, 11). However, not all societies followed the same trajectory. Some remain cooperative only at a smaller, subnational scale (e.g. along ethnic or regional lines), and historically large societies can even backslide toward fragmentation (12–16). Moreover, the overall decline in violence is punctuated by periods of intense conflict, such as the two world wars. The mechanisms underpinning different scales of cooperation (i.e., cooperating only with your close relatives or neighbors vs. cooperating with millions of unknown strangers in the same society) are fairly well understood. What remains poorly understood is how societies shift between mechanisms (e.g. institutions overcome kin-based clans), or what might incentivize a transition to larger-scale cooperation (17).

One way to quantify differences in cooperative scale is through energy capture and population size, which are tightly correlated (18–21). Consider, for example, the transition from burning wood to mining coal. Gathering wood fuel is feasible for a few individuals and yields modest energy, whereas coal mining demands highly coordinated labor and infrastructure, but provides vastly greater energy returns (22). In general, higher-payoff resources (like coal, oil, or nuclear energy) require more cooperative effort to exploit (23), but also support much larger populations once harnessed (24–27). Each major energy innovation in human history, from agriculture to fossil fuels, required a larger-scale cooperative enterprise, and in turn produced a surplus that enabled further population growth. This interdependence of population and resources suggests a positive feedback loop: cooperation begets resources, which beget population growth, which can beget further cooperation by enabling exploitation of new resources.

We propose that the scale of cooperation in society is fundamentally limited by the scale of the reward that cooperation can achieve. In other words, groups cannot sustainably cooperate beyond what their attainable collective benefits support. We use the Stag Hunt game as a framework to model multiple cooperative equilibria in societies facing multiple potential “stag” opportunities. In the classic Stag Hunt, a solitary hunter can always catch a low value here, but a higher value stag requires two hunters to cooperate (28, 29). Pacheco et al. (30) generalized this to an N-person Stag Hunt, effectively a public-goods dilemma with a threshold number of cooperators required to secure the higher reward, and where everyone shares the reward if the hunt succeeds. We extend this concept to multiple reward levels and introduce population dynamics coupling rewards to carrying capacity. For example, a small group can capture only small rewards (e.g. firewood), whereas only a large, highly coordinated group can capture the biggest “stag” (e.g. harnessing a rich energy source like oil or nuclear power). Crucially, the available rewards come in discrete steps of increasing size, and reaching each larger reward requires crossing a corresponding cooperation threshold. If too few people cooperate, the larger reward is simply unattainable. In this case, the collective effort fails (analogous to a single hunter approaching a stag in the classic game), and individuals are better off sticking to smaller, easier payoffs (analogous to a hare in the classic game). This threshold requirement creates a cooperation barrier. Cooperation will not increase beyond the current level unless the population, and therefore the number of potential cooperators, grows sufficiently to make the next reward level attainable. Conversely, once a new threshold is within reach, the model predicts a surge of cooperation to seize that opportunity, which in turn causes a population boom (thanks to the newfound surplus of food, energy, Haber-Bosch synthetic fertilizer, or other population-supporting resources). However, a population increase does not automatically translate into a higher cooperation rate, because a larger population also creates resource pressure (more mouths to feed from the same resource) which can promote selfish behavior and defection if no new reward has been secured. These dynamics may explain both the general expansion of cooperation over human history and the persistent gaps between societies. For instance, a small-scale society sitting atop a bountiful resource (e.g., a hunter-gatherer band living on coal-rich land) cannot instantly exploit that big “stag” without first growing and organizing itself. They would need to develop the technology and social organization to mine and utilize the coal, which are processes requiring larger scales of cooperation. If this leap in cooperative scale is not achieved, the society may remain stuck at a lower level of resource use, or become vulnerable to that resource eventually being exploited by a larger, more cooperative society (e.g., a multinational corporation from an industrialized nation).

In our model, we take a standard approach from evolutionary game theory and cultural evolution for how to model behavior and population change (6, 31), assuming individuals adopt cooperation if it yields higher payoffs than defection (akin to a replicator dynamic), and societies grow or shrink depending on average payoffs (population dynamics using logistic growth). We also model the effects of technology and limited resources. Technological advancement reduces how many people are needed to reach further thresholds. If population size and technology are related as suggested by previous work (32–34), then as technology advances, fewer cooperators are required to maintain the same level of reward per person. That is, more cooperation is initially required to invent and implement the technology, but as we discover means to be more efficient, fewer cooperators are needed. Resource return decay, such as coal or oil becoming more scarce and difficult to acquire (24), reduce the returns from a reward over time, effectively causing the cooperation ladder to slip down, requiring the group to reach greater rewards to sustain the same level of cooperation. Considering technological advancements and resource return decay moves the equilibria of our model but does not affect the general dynamics. All mathematical details (replicator equations, payoff functions, parameter definitions) are provided in the Supplementary Information, allowing us to focus here on intuitive explanations and broad outcomes.

We first describe the model’s structure in intuitive terms. We then present key insights for how cooperation can surge, stall, or regress. In brief, our model reveals that:

1. **Cooperation surges near thresholds:** Cooperation rates increase sharply whenever a new collective reward threshold is almost attainable (e.g. when just one or few additional cooperator/s would enable the group to reach the next “stag”). It is much easier to recruit that last few percent of cooperators needed for a big payoff.
2. **Cooperation stalls between thresholds:** When the next threshold is far out of reach, cooperation tends to stagnate or even decline. Individuals then have less incentive to cooperate and may revert to defection, knowing that their effort won’t secure a higher reward.
3. **Population and cooperation positively feedback:** There is a strong feedback loop between cooperation and population. Cooperative successes raise the group’s carrying capacity and population, which in turn can enable further cooperation by making higher thresholds attainable.
4. **Potential for trap at suboptimal scale:** This virtuous cycle is not guaranteed. Societies can become trapped in stable equilibria below their cooperative potential, especially if early cooperation falters or if thresholds are too distant. In other words, the mere presence of a high-value resource (a big stag) is not enough if the society’s prior scale of cooperation (and population) isn’t sufficient to approach that threshold. History and initial conditions (e.g. an early burst of cooperation or a lack thereof) can determine which equilibrium a society settles in.
5. **Technology lowers cooperative requirements:** Technological advancement reduces the number of cooperators needed to achieve a given payoff, effectively making it easier to climb to higher rungs on the cooperation ladder. This can help groups escape suboptimal equilibria if they can grow enough to develop and adopt the technology. However, in the long run, extremely efficient technology can also reduce the equilibrium proportion of cooperators needed (since tasks become easier, a group can succeed with fewer cooperators).
6. **Resource limits raise cooperative requirements:** Conversely, diminishing returns or resource depletion means that as population grows, the per-capita payoff from the current resource drops, creating a pressure for the group to seek an even higher reward just to maintain cooperation. Finite resources thus exacerbate the need for continual cooperative innovation (climbing to the next threshold).

Together, these results provide a mechanistic, materialist account of how large-scale cooperation emerges, sometimes stalls, and can even regress. In the following, we present our model and results in detail, then discuss implications for cultural evolution and modern collective action problems, highlighting, for example, how technology and resource constraints interact in our model, and how a plateau in resource returns could jeopardize future global cooperation.

## Model

To investigate how the scale of cooperation expands or declines, we build on Pacheco et al.’s generalization of a classic stag hunt game to an *N*-person stag hunt, further generalizing the model by introducing multiple stags requiring different numbers of cooperators, and population growth dynamics tied to cooperative rewards through carrying capacity. We also consider technological advancement and changing resource limits.

Let’s start with Pacheco et al.’s model with a single stag and multiple players but introduce variable cooperative thresholds and rewards. So instead of requiring two hunters, consider a threshold of *M* hunters required to catch a stag. Players can either cooperate and participate in the communal stag hunt, or defect and hunt for a hare on their own. Cooperators thus pay a cost-*c* for cooperating by forgoing the guaranteed reward of a hare. If the number of cooperators *k* is less than the threshold (*k* < *M*), then cooperators gain nothing because the hunt failed and pay the cooperation cost-*c*. For the stag hunt to be successful we require *k* ≥ *M* cooperators. If the hunt succeeds then all members of the group benefit, even those who defected receive their share of the stag as well as the hare they caught, creating a cooperative dilemma. The stag provides *F* food which gets divided equally amongst all group members, providing everyone 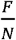 food.

We further generalize this dilemma in four critical ways:

1. **Multiple escalating thresholds:** Instead of a single stag and corresponding threshold, our societies face multiple possible “stags” of exponentially increasing size. Each larger reward demands a larger number of people to cooperate. This reflects real-world patterns such as for energy capture. Higher Energy Return on Investment (EROI) sources (e.g. coal, oil, nuclear) offer much greater payoffs but require larger-scale operations (e.g. extraction, processing, pipelines, protecting, financing, and so on) to exploit (24).
2. **Population dynamics:** The total population is not fixed. It grows when cooperation yields surplus resources (increasing the environment’s carrying capacity), and it declines when resources become scarce. In effect, successful cooperation raises the carrying capacity (more sustenance can support more people), whereas a growing population sharing a fixed resource lowers per capita returns. This introduces a feedback loop between cooperation and population, where cooperation can fuel population growth, and a larger population in turn changes the stakes for cooperation.
3. **Technological advancement:** We consider the effect of technological advancement, which reduces the number of required cooperators needed for a cooperative success. This reflects that as groups increase in size they develop tools and technologies which make cooperation easier (34). In the model, technology effectively makes it easier to reach higher rewards at a given population size. When the technology parameter is set to zero, the model has no technological effects.
4. **Resource limitations:** We also consider finite resource effects, which reduce the benefits of cooperation as population grows. Initially, groups exploit easy, high-yield resources; as those are exhausted, additional people must turn to resources that require more effort or are less abundant, reducing the average benefit of cooperation. This factor has the effect of capping unbounded growth and represents environmental constraints on payoffs.

These extensions allow us to analyze when and how the scale of cooperation in a society increases or decreases. To illustrate the intuition, consider a small group of cooperators that routinely manage to catch a stag (the first threshold). This successful group enjoys high returns with more food and energy than the hunters immediately need. As a result, the group will be able to sustain a larger population over time. With a larger population, the group can tackle a bigger challenge than a stag. For instance, there might now be enough hunters to organize a bison hunt. With sufficient cooperators in the larger group, a bison hunt can succeed, yielding even more food and energy than the stag did. This surplus supports further population growth, which then increases the scale of possible future hunts. In this way, the group climbs the “cooperation ladder” from hare to stag to bison, and so on. The qualitative analogy extends to human history. Hunter-gatherers mainly harnessed fire (comparable to hares), early agriculturalists systematically tapped solar energy through farming (a larger-scale stag), and modern industrial societies exploit coal, oil, natural gas, and nuclear power (even bigger “bison”-sized payoffs) (20, 23). Each of these transitions was accompanied by an exponential increase in energy return on energy invested (EROI) and supported a leap in population size. However, accessing each new energy source required more cooperators working together than before - larger labor forces, complex coordination, and new institutions or technologies. In our model, the returns to cooperation increase super-linearly with group size, such that each threshold crossed yields a disproportionately larger payoff, but crossing each threshold demands a commensurate increase in cooperative effort.

## Results

### Cooperation spikes at thresholds (and tolerates free-riders)

Our model predicts that cooperation increases dramatically whenever a new payoff threshold, or “stag”, becomes within reach. What counts as within reach depends on model variables and parameters (population size, current technology, etc), but the key pattern is that the rate of change in cooperation is highest when the group is just one cooperator away from reaching a threshold. In other words, the incentive for an individual to become a cooperator peaks when only a few more cooperators are needed to achieve a much larger reward. Conversely, the tendency to cooperate is lowest midway between thresholds, when the next reward is far off and you can instead enjoy your share of the current reward. Figure 1 illustrates this basic intuition using a simplified version of the model (holding population fixed and ignoring technology and resource limits for clarity – full details are given in the Supplementary Information). When the group is one cooperator short of the next threshold, the marginal benefit of contributing (cooperating) is greatest and thus cooperation levels surge. In contrast, between thresholds, adding a cooperator does not unlock a new reward, so individuals have less incentive and the cooperation rate dips. *M*oreover, as thresholds get higher (each threshold representing a reward exponentially larger than the last), the “surge” in cooperation when approaching those thresholds becomes more pronounced. Larger rewards create stronger incentives, if they are attainable.

**Figure 1.**
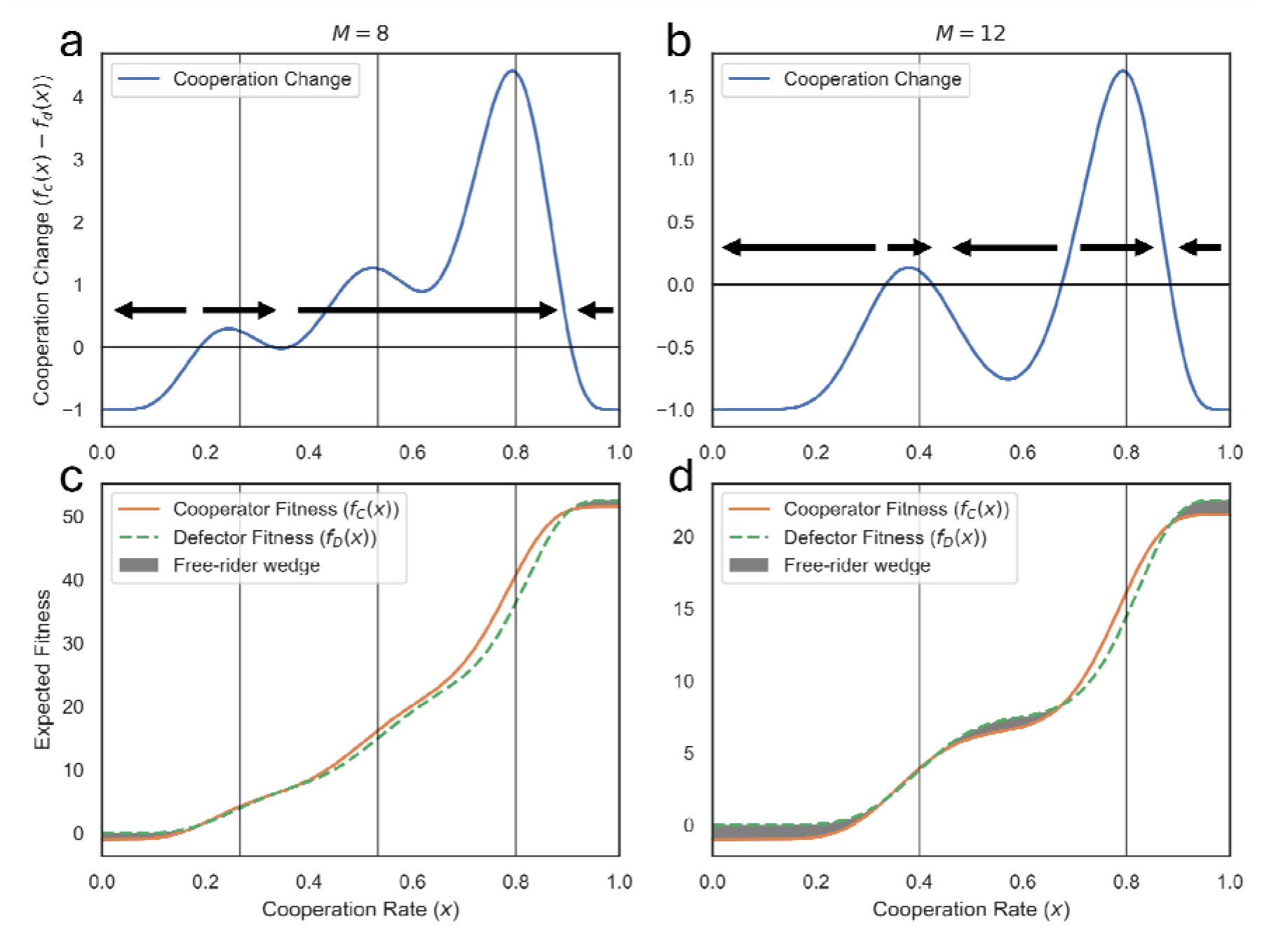
Thresholds Create Surges and Stalls in Cooperation. **Top (a, b):** Show how the current cooperation rate affects its own rate of change. When the curve is above 0, cooperation will increase; below 0, cooperation will decrease; points at 0 are equilibria. Arrows indicate the direction of cooperation change towards equilibria. Notice that cooperation change spikes right before new cooperative thresholds (highlighted by vertical lines). In between thresholds, cooperation rate dips and groups can get stuck at non-optimal equilibria. **Bottom (c, d):** Expected fitness payoffs of cooperators (orange) and defectors (green) with respect to the cooperation rate. This illustrates why the surges, stalls, and reversals occur, with **(c)** corresponding to **(a)** and **(d)** corresponding to **(b)**. The shaded area represents a case where defectors have a greater fitness than cooperators, leading to free-riding and a decrease in cooperation. We see that right before a threshold, both defector and cooperator fitness spike, but cooperator fitness increases faster. In between thresholds, defector fitness overtakes cooperator fitness. (Parameters: group size, cooperation cost, cooperation benefit, threshold multiplier)

An important outcome, which replicates Pacheco et al. (30) is that at any attainted reward level, a certain degree of free-riding is tolerable without collapsing cooperation. Once a threshold reward is secured, even though defectors individually do better than cooperators (since they get the reward without paying the cost), cooperators have no incentive to defect as long as doing so would cause the group to fall below the threshold. In other words, if dropping one cooperator would forfeit the large reward, cooperators cooperate despite the free-riders. Our model reveals a further insight - free-riding can also be stable before the higher thresholds are reached. One might expect that selfish behavior would only be tolerated once the group has achieved the highest threshold reward (i.e. once a pre-industrial society industrializes by exploiting their resources). However, we find that a society can stabilize at an intermediate level of cooperation where some individuals defect even though a higher collective payoff would be possible in theory. In these cases, defectors are not only getting a higher payoff than the cooperators at the current level, but they are also collectively limiting the group’s potential to reach the next threshold (which would benefit everyone, including those defectors). Paradoxically, even though a bigger reward exists, it remains effectively out of reach, making cooperation towards it seem futile for any one individual. This result reveals how societies can become stuck in globally suboptimal equilibrium “kludges”, such as corruption leading to lower payoffs.

### Population booms follow cooperative breakthroughs

When we allow population to change endogenously, the model reveals a powerful feedback loop. Successful cooperation leads to population growth, which in turn can make further cooperation easier by providing more potential cooperators. We first examine this by holding the cooperation rate fixed and analyzing the population dynamics. As shown in Figure 2, the maximum sustainable population (carrying capacity) of the society jumps upward right after a new reward threshold is reached. Between thresholds, however, carrying capacity actually declines slightly with increasing population, because if the group hasn’t yet unlocked a new resource, each additional person just means the same fixed pie is split among more individuals. In Figure 2a, vertical lines mark the thresholds; note the slight dip in carrying capacity between them. The diagonal line indicates where the current population equals the carrying capacity; whenever the population exceeds carrying capacity, population growth becomes negative (the population will decline back toward the capacity). Figure 2b plots the actual population growth rate as a function of population size, showing that growth spikes after each threshold (when resources per capita suddenly jump) and then dips below zero if population overshoots the available resources.

**Figure 2.**
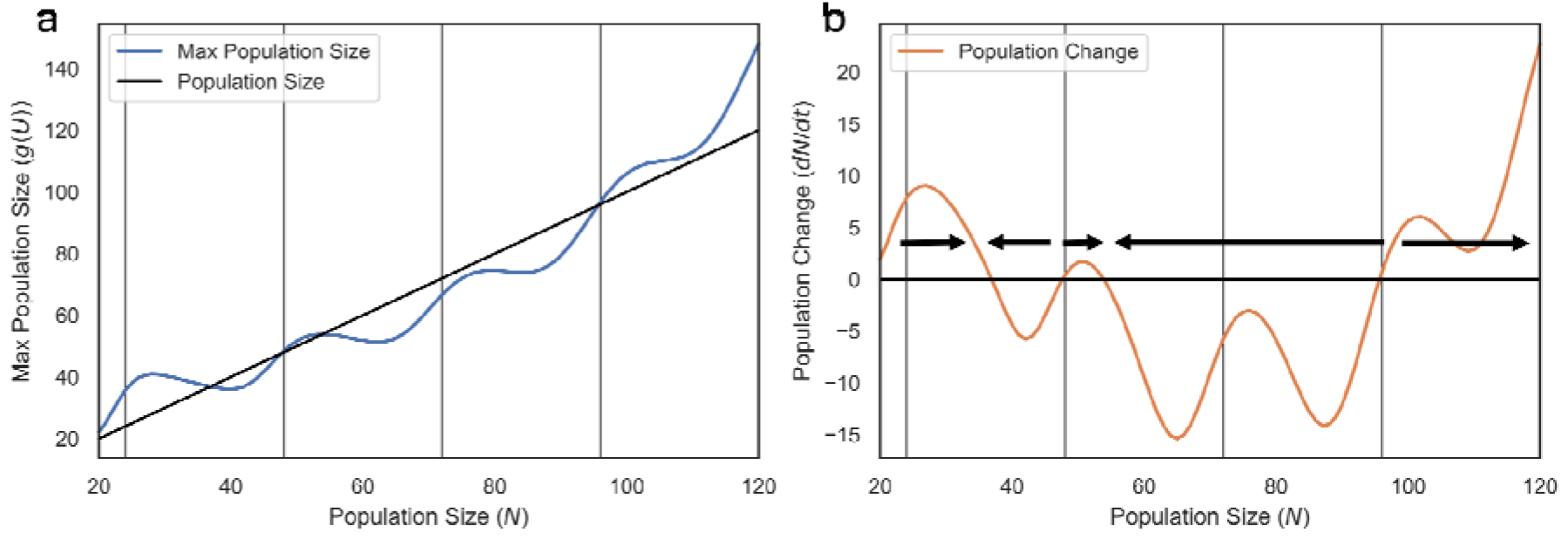
Population Spikes After Reaching Threshold. The effect of population size () on maximum population size **(a)** and population growth **(b). (a)** The maximum population size a group can sustain spikes right after a threshold is reached, with thresholds being marked by vertical lines. In between thresholds, the maximum population slightly decreases as a larger N means the same reward is divided among more individuals. Notice that the current population (evidenced by the diagonal line) can be larger than the maximum population and in this case population growth is negative. **(b)** Population growth similarly spikes after thresholds (vertical lines) and dips in between thresholds. Current population being greater than the maximum population in **(a)** represents a decline in population in **(b)**. This shows that even though for large enough populations, population growth may be positive, the group can get ‘stuck’ at a lower population unable to reach further thresholds. (Parameters: cooperation rate, cooperation cost, cooperation benefit, threshold multiplier, threshold size, baseline population size)

One result visible in Figure 2b is that beyond a certain threshold, the model (without resource constraints) would allow the population to grow without bound - essentially a runaway positive feedback. If each cooperative leap automatically puts the group at the brink of the next threshold, a cascade of breakthroughs could lead to unbounded growth. In reality, available energy and resources are finite. To model this, we include resource decay. As the population grows, the returns from cooperation begin to diminish (e.g. resources become harder to extract). More specifically, we capture resource limitations with per-capita benefit of a given reward decreasing as population increases (our finite-resource parameter). Figure 3 demonstrates how even a modest resource saturation effect can cap the runaway growth. With no resource constraints (blue curve), once a high enough population is reached, the model would predict indefinite growth. But with even a moderate resource return decay (orange curve), an equilibrium population is reached instead of infinity. In short, environmental limits reintroduce a total carrying capacity for the system even in the presence of exponentially growing cooperative rewards.

**Figure 3.**
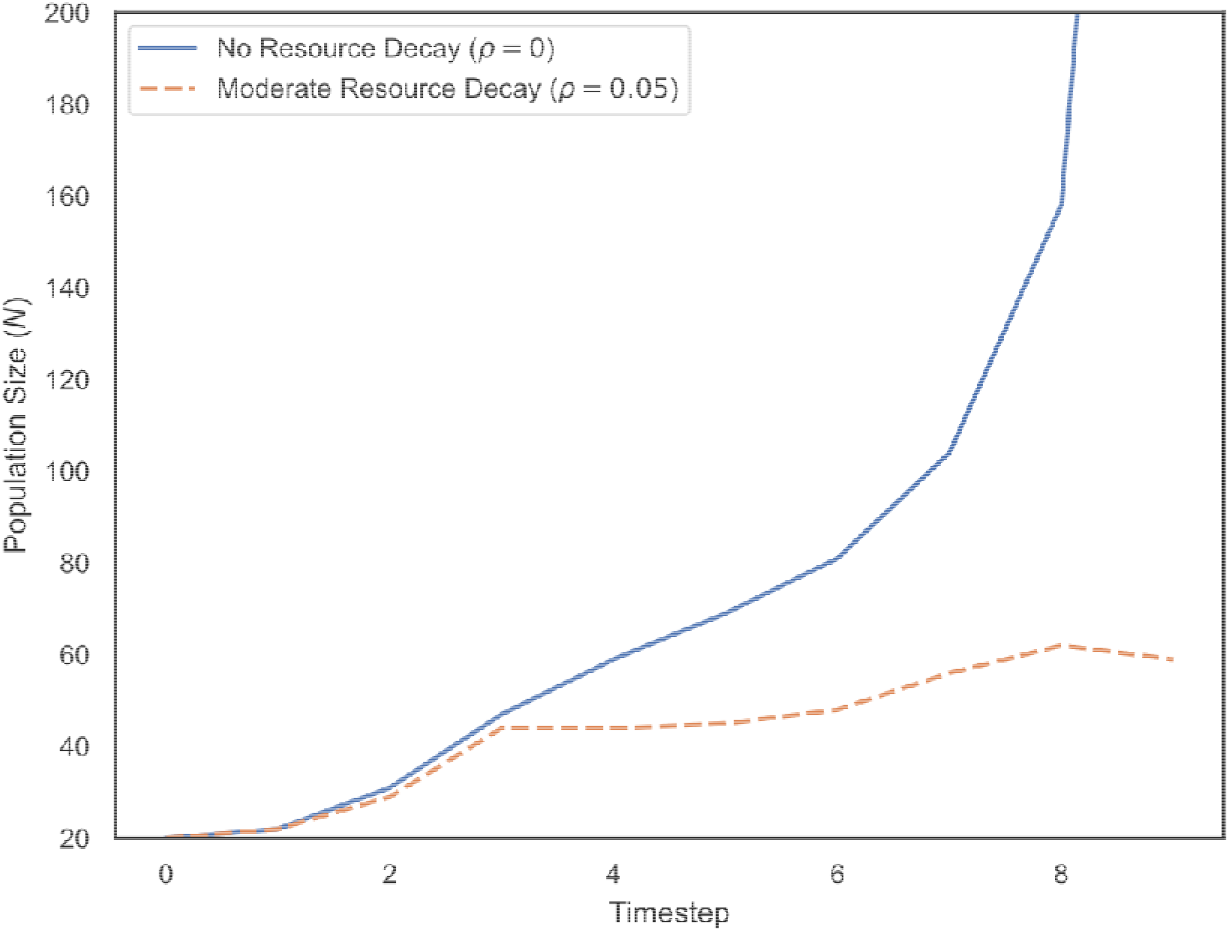
Resource Decay Limits Population Growth. This graph shows population growth over time, for different resource return decays (). When (in blue), then we have the model without any resource constraints, recalling that resource return decay limits the returns from cooperation as population grows. In this case, once the population grows large enough, they can continue to grow indefinitely. However, considering even a moderate decay on cooperative returns with population growth () can act as a cap on population, preventing infinite growth. (Parameters: cooperation rate, cooperation cost, cooperation benefit, threshold multiplier, threshold size, baseline population size, utility strength moderator, population growth rate)

### Feedback between cooperation and population dynamics creates ratcheting (or stagnation)

When we allow both cooperation and population to change together, it reveals a feedback loop between these processes. Near the cooperation threshold for the next reward level, cooperation rate increases pushing the group into a new cooperative scale. This new scale provides high rewards which leads to a population boom and makes a new threshold within reach. The new threshold being within reach means that the cooperation rate will once more increase and the cycle continues. When this feedback loop works then the society can grow and advance. Cooperation levels stay high, payoffs grow, and population size increases without placing a too strong toll on society. However, this virtuous cycle is not guaranteed. Different initial conditions or early outcomes can lead to very different long-term equilibria.

Figure 4 illustrates this path dependence by plotting stable cooperation equilibria at various population sizes. In one scenario (orange trajectory), the group starts with a relatively high initial cooperation level and manages to hit an early threshold, which sets off the ratchet mechanism described above. As population grows, each new threshold is met in turn, and cooperation remains near its peak. The society continually climbs the cooperation ladder. In a second scenario (blue trajectory), the group begins with a lower initial cooperation rate and fails to reach the first threshold. The group remains between thresholds. Instead of striving for a big payoff that seems unattainable, some cooperators gradually defect to recoup their losses (since a larger population sharing the same small reward lowers everyone’s payoff). Cooperation actually declines, and the society settles into a lower-level equilibrium without ever achieving a major breakthrough. These two outcomes emerge despite identical fundamental parameters. The only difference is the initial fraction of cooperators. Thus, small initial advantages (like a slightly higher tendency to cooperate due to cultural pre-adaptations, such as the need to cooperate for defense or agriculture, or a minor technological boost that makes cooperation easier) can compound over time into vastly different societal trajectories. This path-dependent dynamic is analogous to the idea of critical thresholds or “tipping points” in cultural evolution and social change (35).

**Figure 4.**
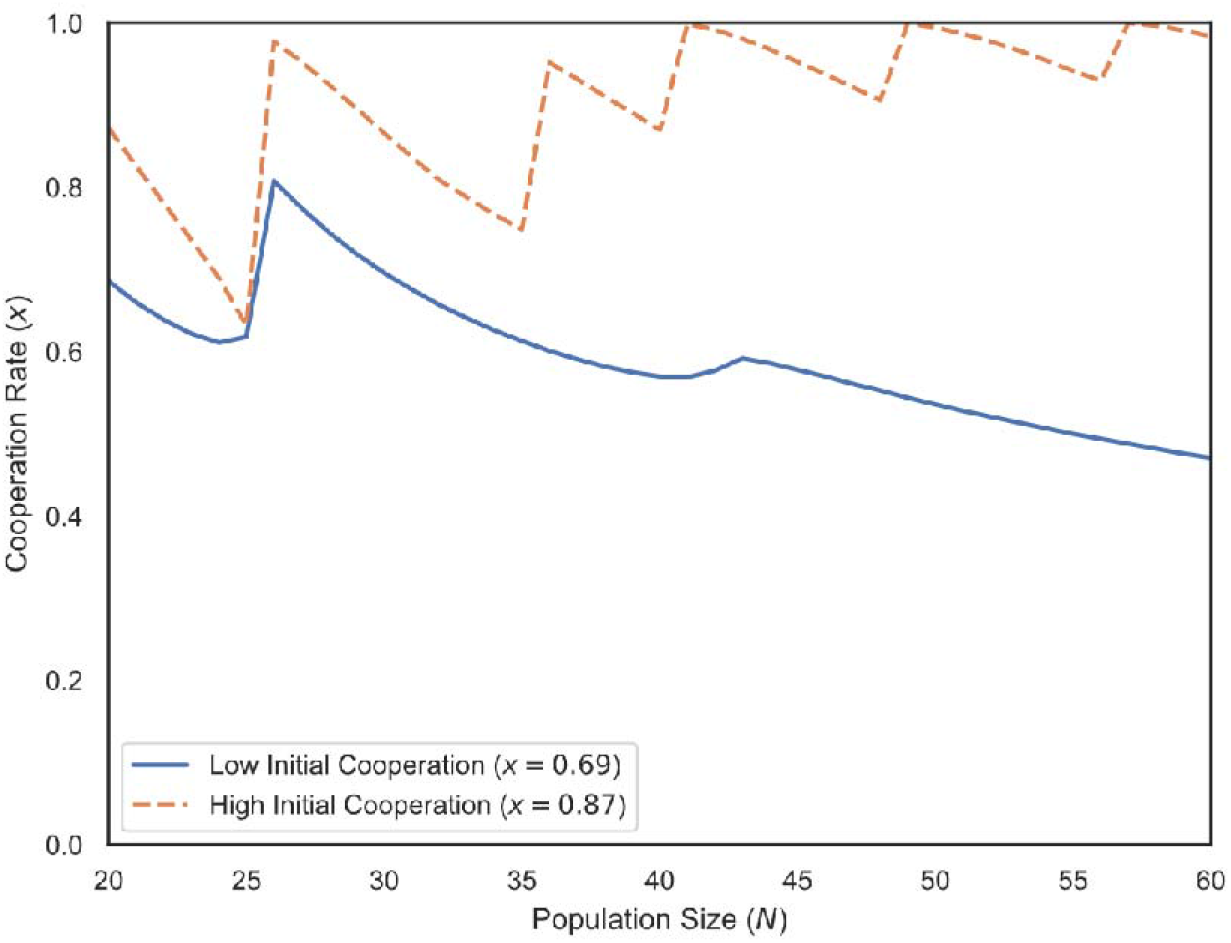
Path Dependencies in Cooperation Rate. Evolutionarily stable cooperation rates as functions of population size under two different starting conditions. Here we see two different equilibrium behaviors dependent on initial cooperation rate. The orange path represents a society that begins with high initial cooperation and thus capitalizes on the positive feedback loop: as population increases, new thresholds keep coming within reach, and cooperation ratchets upward to attain them. The blue path represents a society with a lower initial cooperation rate. As population grows, the group remains stuck below the next threshold, and in fact some cooperators defect due to reduced per capita returns, causing cooperation to decline to a lower equilibrium. Both scenarios assume the same underlying parameters; only the initial state differs. This demonstrates how historical contingency (e.g. an early successful collective action or, conversely, early mistrust) can lock a society into a high-cooperation growth trajectory or a low-cooperation stagnation. (Parameters: cooperation cost, cooperation benefit, threshold multiplier, threshold size, baseline population size, utility strength moderator)

Figure 5 provides a phase-plane view of the coupled population–cooperation dynamics, summarizing the fate of societies from different starting conditions. Depending on initial population size and cooperation rate, a group can end up in one of several regimes: (i) a low-population, all-defection state (lower left, if they start very small and uncooperative); (ii) an intermediate “stuck” state with some cooperation but below the highest potential (middle area, a stable local equilibrium); or (iii) a high-population, high-cooperation trajectory that effectively keeps climbing the ladder (upper right, where each threshold reached leads to the next). A key insight from this figure is that if the initial population is too low, even a high willingness to cooperate might not allow the group to ever reach the first threshold – there is a minimum critical mass needed to get started. Moreover, once on a trajectory, it can be hard to switch paths; the equilibria act like attractors. Notably, if initial cooperation is zero (no one cooperates at first), the group will remain stuck in the all-defector state. This underscores the importance of early cooperative mechanisms (e.g. kin-based, reciprocity, or reputation-based cooperation at small scale) as a seed for larger-scale cooperation to grow.

**Figure 5.**
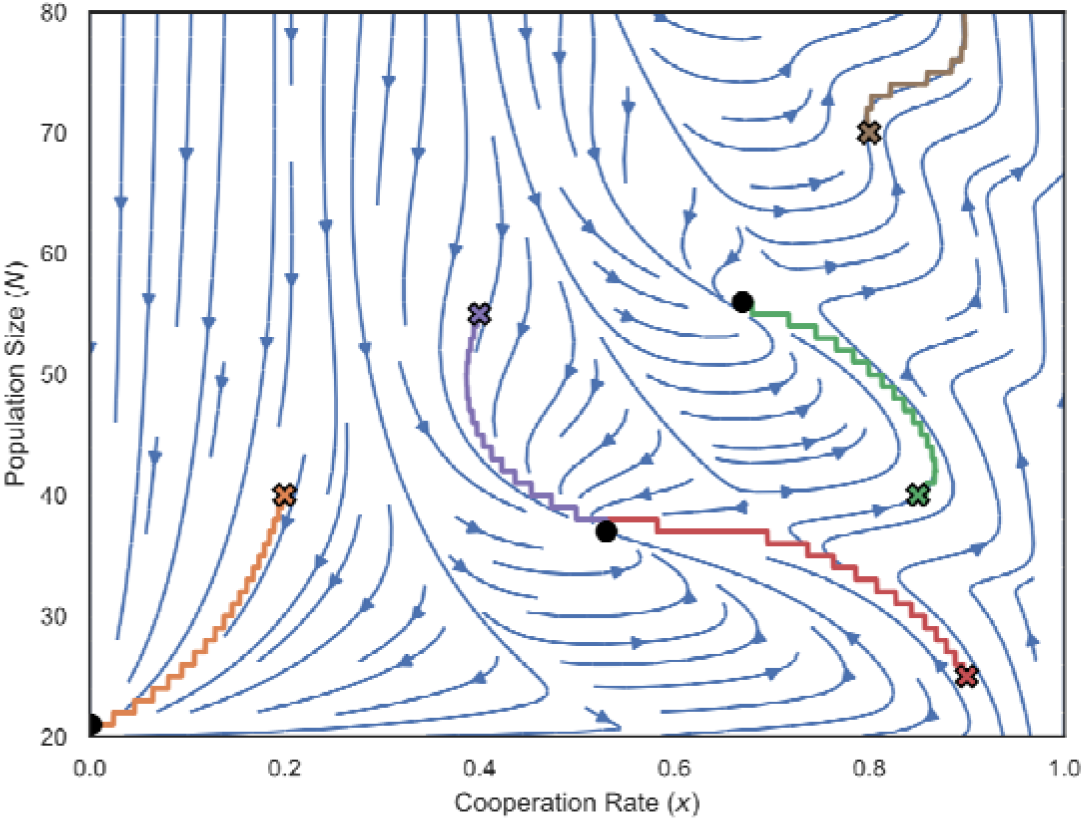
Equilibrium Dynamics of Cooperation Rate () and Population Size () Phase plot of cooperation rate () and population Size (), with example starting cooperation rate and population size pairs (marked by Xs), along with their trajectories. We generally see that the phase plot is split into quadrants for a corresponding equilibrium condition (marked by dots). When cooperation rate is quite low then the group falls into the all defection and low population condition. Higher cooperation corresponds to a more rewarding internal equilibrium. If the cooperation rate and population size are sufficiently high, then the group will be able to keep reaching new thresholds and continue to climb in population size. We also see that if population is too low, then even a very high cooperation rate may not allow the group to reach a better equilibrium. The number and location of equilibria is dependent on the model parameters. (Parameters: cooperation cost, cooperation benefit, threshold multiplier, threshold size, baseline population size, utility strength moderator)

### Effects of technology and resource constraints

Technological advancement and resource depletion modulate these dynamics in intuitive ways. Technology, by lowering the effective threshold needed for success, can pull a group out of a suboptimal equilibrium. In our model we tie the technology level to population size (consistent with literature suggesting larger populations tend to generate or sustain more complex technologies (33). As the society grows, technology makes cooperation incrementally easier, helping prevent long plateaus. However, in terms of ultimate equilibrium, a high level of technology can actually result in a lower fraction of cooperators needed at equilibrium – essentially because tasks become so efficient that not everyone needs to contribute. Figure 6 illustrates that as the technology factor increases, the stable cooperation rate at equilibrium can decrease (since a smaller cooperative effort achieves the same large reward). On the other hand, introducing a stronger resource decay (finite resource effect) has the opposite effect, increasing the required cooperation level at equilibrium. When each person’s contribution yields less (due to resource scarcity), successful collective action becomes more crucial to maintain welfare. Importantly, in our model these factors change some details of the transient and final dynamics but do not alter the overall pattern of surges and stalls. A sufficiently high technology can make the “infinite growth” outcome (upper right of Figure 5) more likely by smoothing the path between thresholds, whereas resource limitations push more societies into needing continuous innovation (higher thresholds) to avoid stagnation.

**Figure 6.**
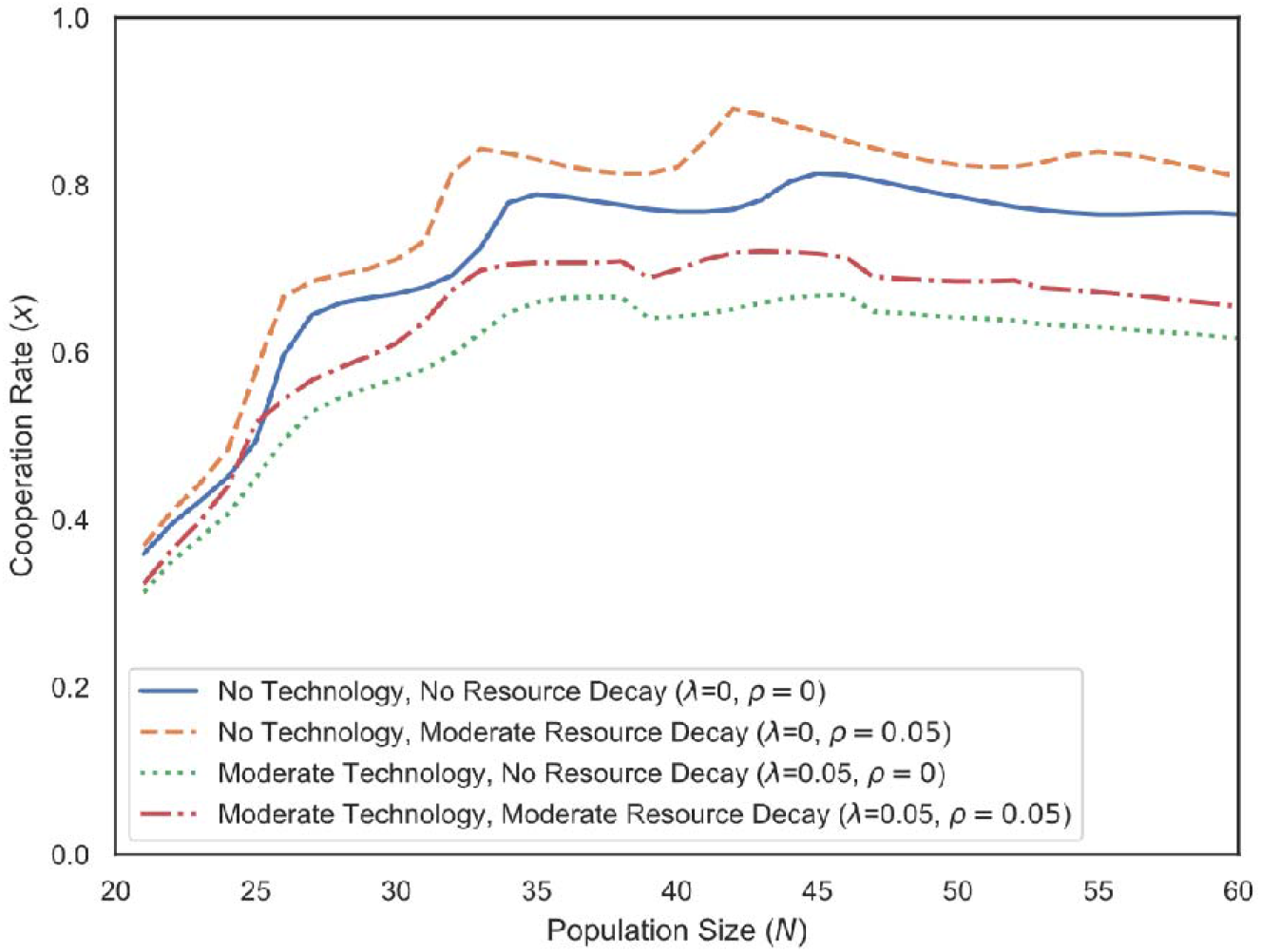
Technology Eases Pressure to Cooperate, Resource Decay Increases Pressure to Cooperate. Equilibrium cooperation rates and population sizes as a function of technological efficiency and resource scarcity. Higher technology (moving rightward) reduces the fraction of cooperators needed to achieve a given output, thus the stable cooperation rate declines – the society can function with more free-riders because cooperation is easier. In contrast, greater resource decay (moving upward) means per capita returns drop faster with population, so maintaining a given living standard requires more cooperation; thus stable cooperation rates increase. (For this figure, only internal equilibrium states are shown; extremely high population cases leading to unbounded growth are not depicted. Parameters as before, with varying technology and resource-decay factors). (Parameters: cooperation cost, cooperation benefit, threshold multiplier, threshold size, population size, utility strength moderator)

## Discussion

We introduce a theoretical model to explore how human cooperation can scale up in tandem with population growth and material payoffs. By generalizing the *N*-person Stag Hunt to allow multiple escalating thresholds we capture a multi-stage cooperation dynamic. When a new cooperative opportunity becomes viable, the incentive to cooperate sharply rises. But if no higher goal is attainable, individuals tend to revert to competition or tolerate free-riding. Incorporating population dynamics, technology, and finite resources makes these swings even more pronounced. A growing population can enable larger cooperative endeavors but also imposes resource pressure. If that pressure is not relieved by reaching the next threshold, cooperation can temporarily erode until the group either collapses or stabilizes at a lower level. The result is cooperation ladder, where cooperation climbs in abrupt jumps from one rung (threshold) to the next, and only by climbing does the next rung come within reach. Importantly, our model only explores how existing cooperative groups increase in scale of cooperation, not how cooperation emerges initially nor the specific mechanisms that may be favored at different levels. Both the initial conditions and mechanisms at each scale can be explained by the large body of literature on mechanisms of cooperation (6). For example, kin and reciprocity-based cooperation, as found in other non-human animals may be sufficient for smaller scales, but are insufficient, which may require evolving institutions that are unnecessary for smaller scales. These insights align with and provide a concrete mechanism for several patterns noted in anthropology, history, and economics.

The model helps explain how large-scale cooperation can gradually emerge through intermediate steps. Human societies did not leap from small bands to vast empires overnight; they typically expanded through intermediate scales (e.g., tribes, chiefdoms, states) often associated with new technologies or organizational breakthroughs (e.g., agriculture, city formation, industrialization) (36). In our model, each of these transitions corresponds to crossing a cooperative threshold that yielded a major increase in resources and supporting population growth. This feedback loop between cooperation and population growth model how necessity (population pressure) spurs cooperation, and cooperation (better resources) spurs further growth. We showed how this feedback loop can produce accelerating gains and why this process can stall or even reverse in some cases. Not every group in history managed to unlock the next reward level. Those that did, perhaps through favorable geography or chance inventions, surged ahead in scale, while others remained relatively small. In some cases, interaction with other large cooperative groups may have further hindered initial conditions as Nunn and Wantchekon (37) have argued for the effect of the transatlantic and Indian ocean slave trade’s long run effects on trust in Africa. Crucially, our model suggests that the absence of large-scale cooperation in some societies is not because those people are less cooperative by nature, but because the initial or material conditions, and thus payoffs, did not support climbing the cooperation ladder (38). This is also consistent with evidence that even very small-scale societies can achieve impressively large-scale cooperation for certain endeavors when needed, such as for intergroup conflict (39–41). These may be instances where a particular goal (like defense or warfare) offered a sufficiently high payoff or existential necessity to effectively raise the incentive for large-scale cooperation, allowing a usually small group to momentarily act “big.” In everyday life, however, those same groups do not routinely cooperate at large scale because the returns of doing so (in mundane tasks) don’t justify the effort.

Our results highlight that the emergence of large-scale cooperation cannot be taken for granted nor assumed to be an automatic one-way trend. Some theories attribute the decline of violence and rise of large societies to ideological progress or normative shifts that “tamed” or “civilized” societies (e.g. the argument that Enlightenment-era humanism or other moral insights made us more cooperative (11)). Our model implies that if the underlying payoff landscape changes, cooperation can recede, even if cultural values remain pro-social. If the group cannot achieve further growth in cooperative returns, then even rational, self-interested cooperators will be incentivized to defect (42). A pertinent modern implication concerns energy and sustainability. If our current global society fails to find new high-EROI energy sources or other means to dramatically increase the payoff for cooperation, we may face a plateau or even a drop in large-scale cooperation. Today’s renewable energy sources generally have lower EROI than past fossil fuels (24, 43). Even with improvements, they may not on their own provide the kind of exponentially increasing energy surplus that our model indicates helped fuel past growth (44). Humanity may thus need an alternative energy breakthrough with vastly larger returns, for example, cheaper nuclear fission or even fusion power, to sustain the current trajectory of growth without increased competition over resources (45). If such a breakthrough (cheaper nuclear, fusion, or otherwise) arrives, it could alleviate resource competition and drive a new wave of cooperative growth (46). But our model offers a caution, that even a major new energy source only “kicks the can down the road”, though the potential of these technologies may kick the can down the road for centuries. Nonetheless, the overall dynamic, such that eventually the ladder ends and resource limits start to bite. This perspective aligns with work on climate change and global commons problems. Solving the climate crisis requires unprecedented cooperation, yet if nations perceive diminishing returns or a zero-sum scenario in energy and resources, they may default to short-term self-interest (47, 48). Our findings underscore that material incentives are fundamental drivers of sustained cooperation. *M*oral appeals or ideological shifts may help transition between levels where there is misperception or even be a social technology for sustaining cooperation, but there is an underlying material reality that groups are cooperating toward. Without that cooperative reward, these norms may be unlikely to sustain cooperation.

There are several limitations of the present model that could be explored in future work. For tractability, we assumed that population growth benefits are immediate and that resources are shared evenly among individuals. In reality, population growth has a time lag (children take years to become productive contributors), which could create short-term strains: a sudden population boom means more mouths to feed now without immediate additional cooperators, potentially inducing a temporary crisis. Incorporating such delays would likely amplify the oscillations we predict. A boom could be followed by a dip in cooperation before the new generation fully contributes. We also assumed equal sharing of cooperative returns, whereas real societies have inequality. Unequal sharing could allow a subset of cooperators to enjoy high payoffs while others benefit little, which might discourage those who receive less from cooperating. Inequality could thus raise the baseline incentive required to start the ratcheting process (since if rewards aren’t broadly shared, the average cooperator’s benefit is lower). However, as long as collective returns grow exponentially with scale, even those getting a smaller slice could still benefit enough to continue cooperating. We expect the core mechanism of our model, with thresholds and feedback loops, to be robust, but the exact conditions for the virtuous cycle might become more stringent under inequality or may need to be analyzed as groups within groups. A long these lines, another aspect for future exploration is inter-group competition. We did not explicitly model warfare or external threats, but historically, competition between groups can spur greater internal cooperation up to a point (49). Our framework could be extended by considering multiple competing societies each trying to reach the next stag first or preventing rivals from doing so.

In summary, we present a theoretical model that provides a materialist, cultural evolutionary explanation for the expansion and limits of human cooperation. Our model shows how cooperation can expand from a small scale to anonymous strangers, how free-riding can persist without collapsing cooperation, and how cooperative regimes can backslide even after long periods of success. The key to understanding these puzzles is in the coevolution of cooperation with population and resources. Societies that could muster enough cooperation at critical junctures, including in initial conditions, reaped outsized benefits (more energy, more people), setting off a self-reinforcing cycle of growth. Those that could not remained at smaller scales, not necessarily due to any ideological failings or lesser “moral” development, but because the initial or material conditions did not favor scaling up to the next level. The framework situates the literature on the evolution of cooperation and the various identified mechanisms within a broader coevolution of culture and ecology. The model also offers insights into how we may foster global cooperation. For challenges like climate change, this could mean aligning economic and technological incentives such that the next threshold of sustainable energy or collective benefit is visibly worthwhile and within reach, such as through aspirational rather than negatively framed approaches to planetary futures (50).

## Materials and Methods

Table 1 summarizes the key parameters of the model. Table 2 summarizes the key state variables.

**Table 1:**
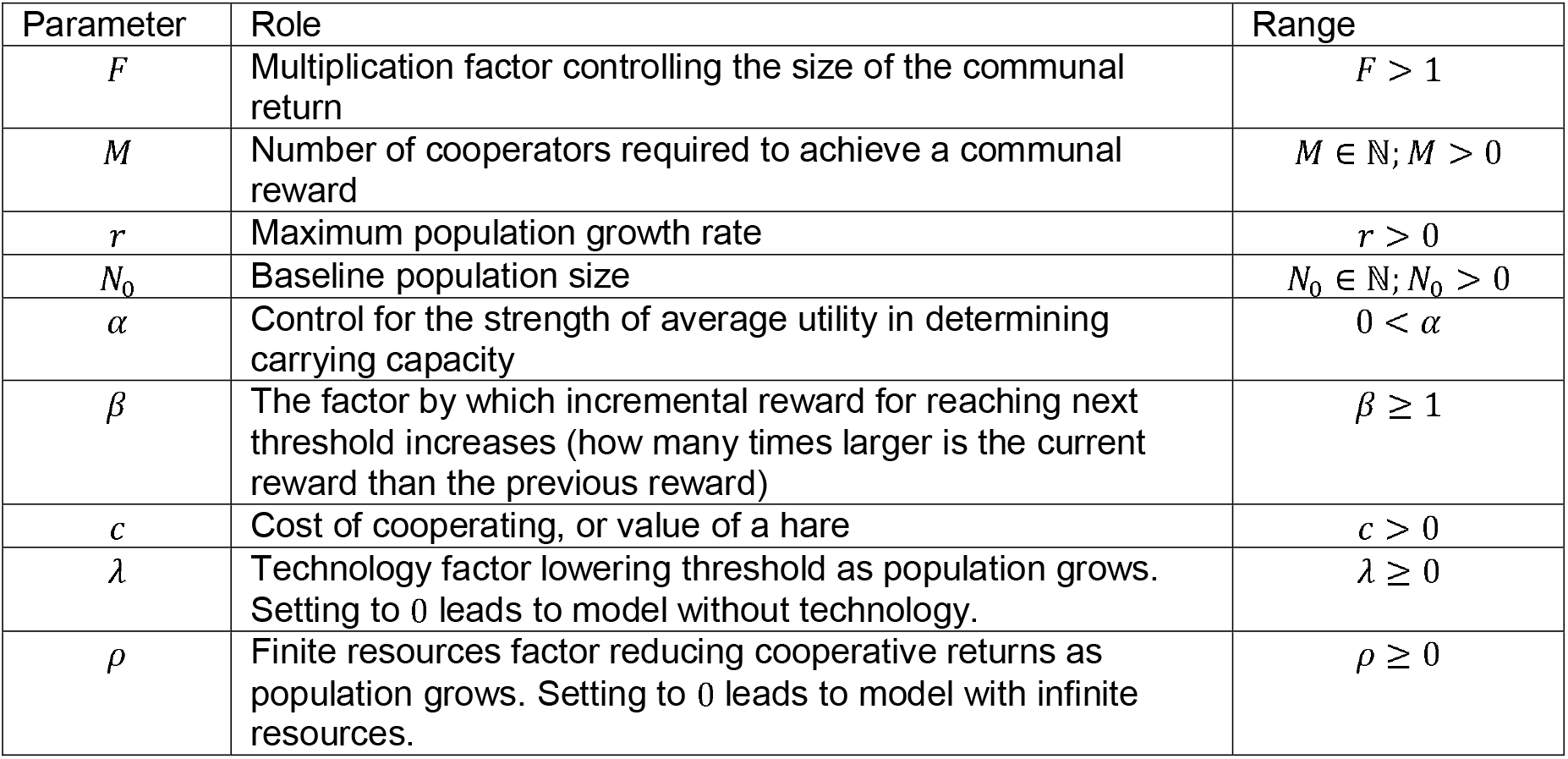
Parameters of the Model.

**Table 2:**
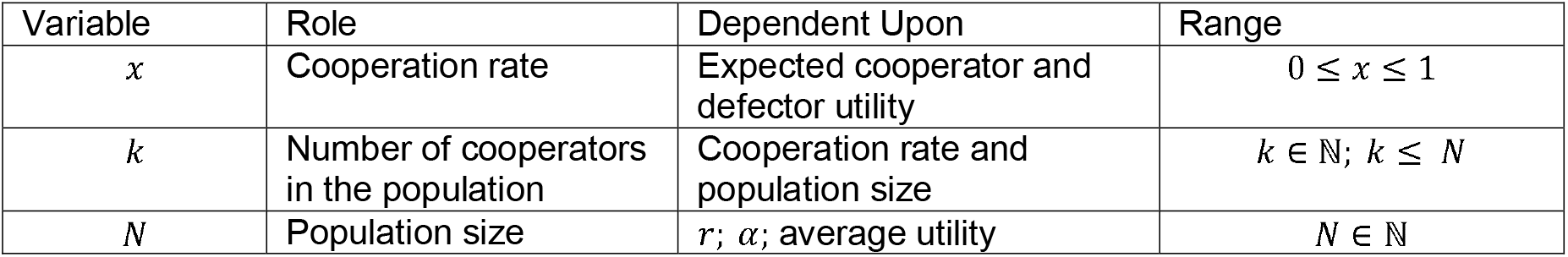
Variables of the Model.

### Model Framework

The utility functions of defectors (*U*_*D*_) and cooperators (*U*_*C*_) are given by Equations 1 and 2:

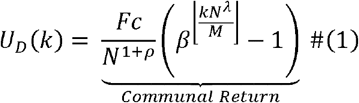

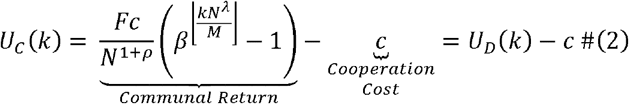

Equations (1) and (2) share a common portion, 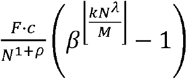, which we call the communal return. Every member of the community receives this return. However, as shown in equation (2), cooperators also pay an additional cost *c* to their utility. By cooperating in the stag hunt, cooperators miss the opportunity for an individual reward and thus pay a cost of *c*.

The communal return is in two parts: 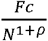 and 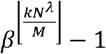. This first term represents the utility of the communal return one player gets in relation to the utility they would get from defecting. In other words, the cooperative reward is times larger than the cooperation cost *c*. Because of the resource decay factor (*ρ*), cooperative returns decrease as population increases. This is meant to capture that resource extraction gets more difficult with time as easily accessible resources are consumed first with more difficult resources consumed later (23). We use population size as a proxy of time. The second term is a multiplicative factor on the communal return, determined by which level of cooperation the group is currently at. For simplicity, let’s assume λ = 0 so we are left with 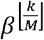. Thus, *M* is the threshold for the number of cooperators required to succeed at a hunt. For every *M* cooperators the next level of communal return is reached, with the size of the return increasing for each reached threshold. *β* represents how much larger the communal return gets for each new threshold reached. So, for every *M* cooperators, the size of the communal return grows exponentially by some rate *β*. This dynamic is achieved in the floor function 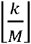. Note that the increase in communal return doesn’t necessarily need to be exponential, but rather could be any stepwise increasing function, with relation to 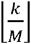. For the purpose of our model we use an exponential function as it most accurately reflects the real world increases in EROI across technological innovation (20, 24). The minus 1 in this term is there to ensure that if the lowest threshold is not met then there is no communal return.

### Evolutionary Dynamics

The likelihood of any given individual in the population cooperating is given by the cooperation rate *x* and we study its evolution using replicator dynamics, as specified in (3):

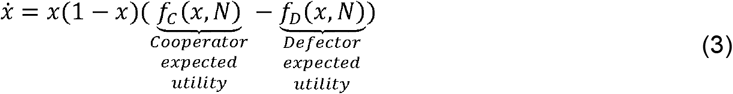

Equation (3) is dependent on expected utility of cooperators and defectors, which are themselves specified below:

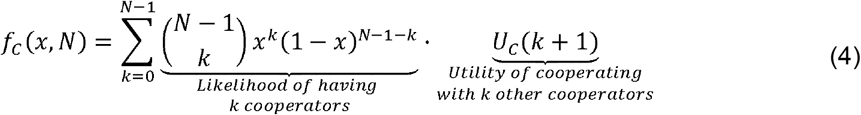

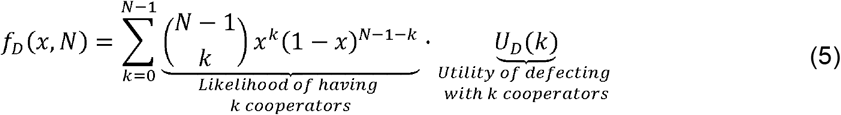

In essence, Equation (4) tells the expected utility of cooperating, calculated as the likelihood of there being *k* cooperators out of the other *N* – 1 members of the group and what utility a player would have by becoming the *k* + 1 cooperator. Equation (5) is the expected utility of defecting, from the likelihood of there being *k* cooperators out of the other *N* – 1 members and the player’s utility if they chose to defect. In practice this works as sampling *N* – 1 people from a well-mixed infinite population with cooperator frequency *x* and defector frequency 1–*x*. This process is standard in evolutionary game theory and lets us study how cooperation rate effects frequency-17 dependent strategy selection without concerning ourselves with problem such as network structure. Also note how *U*_*D*_(*k*) > *U*_*C*_(*k*) for all *k*, meaning that defection is always individually more beneficial than cooperation, however in choosing whether to cooperate, players actually consider *U*_*C*_(*k* + 1) because they are considering the benefits of cooperation while joining an existing group of *k* cooperators. This creates a classic cooperation dilemma, where defectors outcompete cooperators by benefiting from free-riding, however players can still be incentivized to cooperate themselves.

We then calculate population growth modeled as logistic growth in equation (6):

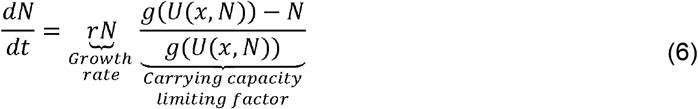

This is dependent on carrying capacity (*g*(*U*(*x,N*))) specified in equation (7), which itself is dependent on average utility specified in equation (8).

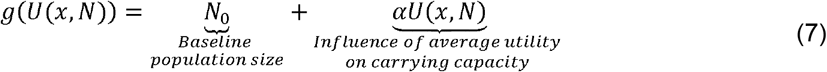

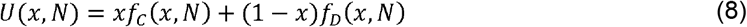

When the current population is below the carrying capacity then it will grow to reach it, when current population is above the carrying capacity then it will shrink to fall back to it. As the utility increases, then so does the carrying capacity. This is because societies which capture more energy can sustain more people (20).

These are the core equations governing the model’s behavior. Full derivations and additional analyses, including solving for equilibrium points, stability, and threshold conditions are provided in the Supplementary Information.

