## Supplemental Information for "The Cooperation Ladder: Scale-dependent payoffs and population dynamics create surges, stalls and reversals"

**Supporting Information Text**

**Parameters and Variables**

Start by recalling our parameters and variables.

| Parameter | Role | Range |
|---|---|---|
| $F$ | Multiplication factor controlling the size of the communal return | $F > 1$ |
| $M$ | Number of cooperators required to achieve a communal reward | $M \in \mathbb{N}; M > 0$ |
| $r$ | Maximum population growth rate | $r > 0$ |
| $N_0$ | Baseline population size | $N_0 \in \mathbb{N}; N_0 > 0$ |
| $\alpha$ | Control for the strength of average utility in determining carrying capacity | $0 < \alpha$ |
| $\beta$ | The factor by which incremental reward for reaching next threshold increases (how many times larger is the current reward than the previous reward) | $\beta \geq 1$ |
| $c$ | Cost of cooperating, or value of a hare | $c > 0$ |
| $\lambda$ | Technology factor lowering threshold as population grows. Setting to $0$ leads to model without technology. | $\lambda \geq 0$ |
| $\rho$ | Finite resources factor reducing cooperative returns as population grows. Setting to $0$ leads to model with infinite resources. | $\rho \geq 0$ |

**Table S1: Parameters of the Model**

| Variable | Role | Dependent Upon | Range |
|---|---|---|---|
| $x$ | Cooperation rate | Expected cooperator and defector utility | $0 \leq x \leq 1$ |
| $k$ | Number of cooperators in the population | Cooperation rate and population size | $k \in \mathbb{N};\ k \leq\ N$ |
| $N$ | Population size | $r$; $\alpha$; average utility | $N \in \mathbb{N}$ |

**Table S2: Variables of the Model**

**Model Framework**

**Individual Utility**

The following are our utility functions as defined in the paper, where $U_D$ is the utility for a defector (1) and $U_C$ is the utility for a cooperator (2).

$$U_D(k, N) = \frac{Fc}{N^{\rho+1}}\left(\beta^{\left\lfloor\frac{kN^\lambda}{M}\right\rfloor} - 1\right) \tag{1}$$

$$U_C(k, N) = U_D(k) - c = \frac{Fc}{N^{\rho+1}}\left(\beta^{\left\lfloor\frac{kN^\lambda}{M}\right\rfloor} - 1\right) - c \tag{2}$$

The explanations for how we construct these equations can be found in the methods of the main manuscript.

**Average Utility**

The following are our average utility functions as defined in the paper, where $f_D$ is the average utility for a defector (3) and $f_C$ is the average utility for a cooperator (4). These are calculated by looking at the likelihood of having a certain number of defectors and cooperators and then taking the utility of a player either defecting or cooperating in that case.

$$f_D(x,N) = \sum_{k=0}^{N-1} \binom{N-1}{k} x^k (1-x)^{N-1-k} U_D(k,N) \tag{3}$$

$$f_C(x,N) = \sum_{k=0}^{N-1} \binom{N-1}{k} x^k (1-x)^{N-1-k} U_C(k+1,N) \tag{4}$$

The groups average utility is thus defined in equation (5).

$$U(x,N) = x f_C(x,N) + (1-x) f_D(x,N) \tag{5}$$

**Cooperation Change**

Cooperation change over time is given by a replicator equation, as defined in equation (6).

$$\dot{x} = x(1-x)\big(f_C(x,N) - f_D(x,N)\big) \tag{6}$$

When cooperators have a greater expected payoff than defectors then $\dot{x} > 0$ cooperation rate will increase. When defectors have a greater expected payoff than cooperators then $\dot{x} < 0$ cooperation rate will increase.

**Population Change**

We calculate cooperation change using a logistic growth function, given by equation (7).

$$\frac{dN}{dt} = rN \frac{g\big(U(x,N)\big) - N}{g\big(U(x,N)\big)} \tag{7}$$

Where this function is dependent on the carrying capacity ($g\big(U(x,N)\big)$) as defined in equation (8), itself dependent on average utility previously defined in equation (5).

$$g\big(U(x,N)\big) = N_0 + \alpha U(x,N) \tag{8}$$

**Analyzing Cooperation Change**

We are concerned with knowing when cooperation rates will increase or decrease in the population, and the local extrema of cooperation rate changes. That is to say we are interested in the sign of our cooperation change equation, as specified in equation (6), and in solving for $x$ such that the derivative of the cooperation rate change equation is zero. Note that the sign of this equation is given solely by the term $f_C(x,N) - f_D(x,N)$. To analyze the behavior of this component, we will first simplify it.

$$f_C(x,N) - f_D(x,N) = \sum_{k=0}^{N-1} \left[\binom{N-1}{k} x^k (1-x)^{N-1-k} U_C(k+1,N)\right] - \sum_{k=0}^{N-1} \left[\binom{N-1}{k} x^k (1-x)^{N-1-k} U_D(k,N)\right]$$

$$f_C(x,N) - f_D(x,N) = \sum_{k=0}^{N-1} \left[\binom{N-1}{k} x^k (1-x)^{N-1-k} \big(U_C(k+1,N) - U_D(k,N)\big)\right]$$

$$f_C(x,N) - f_D(x,N) = \sum_{k=0}^{N-1}\left[\binom{N-1}{k} x^k (1-x)^{N-1-k}\big(U_D(k+1,N) - c - U_D(k,N)\big)\right]$$

$$f_C(x,N) - f_D(x,N) = \sum_{k=0}^{N-1}\left[\binom{N-1}{k} x^k (1-x)^{N-1-k}\big(U_D(k+1,N) - U_D(k,N)\big) - c\binom{N-1}{k} x^k (1-x)^{N-1-k}\right]$$

And with $\sum_{k=0}^{N-1}\binom{N-1}{k} x^k (1-x)^{N-1-k} = 1$, we get:

$$f_C(x,N) - f_D(x,N) = -c + \sum_{k=0}^{N-1}\left[\binom{N-1}{k} x^k (1-x)^{N-1-k}\big(U_D(k+1,N) - U_D(k,N)\big)\right]$$

$$f_C(x,N) - f_D(x,N) = -c + \sum_{k=0}^{N-1}\left[\binom{N-1}{k} x^k (1-x)^{N-1-k}\left(\frac{Fc}{N^{\rho+1}}\beta^{\left\lfloor\frac{(k+1)N^\lambda}{M}\right\rfloor} - \frac{Fc}{N^{\rho+1}}\beta^{\left\lfloor\frac{kN^\lambda}{M}\right\rfloor}\right)\right]$$

$$f_C(x,N) - f_D(x,N) = -c + \frac{Fc}{N^{\rho+1}}\sum_{k=0}^{N-1}\left[\binom{N-1}{k} x^k (1-x)^{N-1-k}\left(\beta^{\left\lfloor\frac{(k+1)N^\lambda}{M}\right\rfloor} - \beta^{\left\lfloor\frac{kN^\lambda}{M}\right\rfloor}\right)\right]$$

$$f_C(x,N) - f_D(x,N) = c\left(-1 + \frac{F}{N^{\rho+1}}\sum_{k=0}^{N-1}\left[\binom{N-1}{k} x^k (1-x)^{N-1-k}\left(\beta^{\left\lfloor\frac{(k+1)N^\lambda}{M}\right\rfloor} - \beta^{\left\lfloor\frac{kN^\lambda}{M}\right\rfloor}\right)\right]\right)$$

From this we define $Q_\lambda(x,N)$ as equation (9):

$$Q_\lambda(x,N) = \sum_{k=0}^{N-1}\left[\binom{N-1}{k} x^k (1-x)^{N-1-k}\left(\beta^{\left\lfloor\frac{(k+1)N^\lambda}{M}\right\rfloor} - \beta^{\left\lfloor\frac{kN^\lambda}{M}\right\rfloor}\right)\right] \tag{9}$$

Meaning,

$$f_C(x,N) - f_D(x,N) = c\left(-1 + \frac{F}{N^{\rho+1}}Q_\lambda(x,N)\right) \tag{10}$$

And,

$$\dot{x} = cx(1-x)\left(-1 + \frac{F}{N^{\rho+1}}Q_\lambda(x,N)\right) \tag{11}$$

When $x = 0$ or $x = 1$ then the cooperation rate is stable, but we are interested in the interior fixed points of the equation, determined by $Q_\lambda(x,N)$. Here we see that cooperation rate will increase when,

$$Q_\lambda(x,N) > \frac{N^{\rho+1}}{F}$$

And it will decrease when,

$$Q_\lambda(x,N) < \frac{N^{\rho+1}}{F}$$

And cooperation rate is fixed for,

$$Q_{\lambda}(x,N)=\frac{N^{\rho+1}}{F}$$

We will not do any deeper analysis on this result as it is very dependent on some parameters, namely $F$. If we increase $F$, that is make the reward for cooperation larger, then the cooperation rate is likely to increase. Similarly taking into account $\rho$, which represents decreased returns from cooperation as population increases, as a way of operationalizing the higher cost of resource extraction over time, then greater resource costs will lead to a more likely decrease in cooperation rate. To better understand this equation, it is more useful to know when cooperation rate change reaches its extrema. We will study this by looking at the derivative for in $x$ of the function determining our interior fixed points and to do so we take the derivative of $f_C(x,N)-f_D(x,N)$ with respect to $x$ and finding when this equals 0. Or,

$$\frac{FcQ'_{\lambda}(x,N)}{N^{\rho+1}}=0$$

$$Q'_{\lambda}(x,N)=0$$

This remains a difficult question to solve, so instead we will analyze a subcase of the question where there is no effect of technology and so $\lambda=0$.

**Subcase: $\lambda=0$**

When there is no effect of technology in our model (or $\lambda=0$), then we can further simplify $Q_0(x,N)$. Noting that for most values of $k$, $\left\lfloor\frac{k}{M}\right\rfloor=\left\lfloor\frac{k+1}{M}\right\rfloor$, and when this equality holds then $\beta^{\left\lfloor\frac{k+1}{M}\right\rfloor}-\beta^{\left\lfloor\frac{k}{M}\right\rfloor}=0$. As such we need only to evaluate the instances where $\left\lfloor\frac{k}{M}\right\rfloor\neq\left\lfloor\frac{k+1}{M}\right\rfloor$, which can be done by substituting $k=lM-1$, such that:

$$\beta^{\left\lfloor\frac{lM}{M}\right\rfloor}-\beta^{\left\lfloor\frac{lM-1}{M}\right\rfloor}=\beta^{l}-\beta^{l-1}=\beta^{l-1}(\beta-1)$$

and we rewrite $Q_0(x,N)$ as:

$$Q_0(x,N)=\sum_{l=1}^{\left\lfloor\frac{N}{M}\right\rfloor}\left[\binom{N-1}{lM-1}x^{lM-1}(1-x)^{N-lM}\beta^{l-1}(\beta-1)\right]$$

We then we tidy the equation:

$$Q_0(x,N)=\frac{(\beta-1)(1-x)^{N}}{\beta x}\sum_{l=1}^{\left\lfloor\frac{N}{M}\right\rfloor}\left[\binom{N-1}{lM-1}\left(\frac{x}{1-x}\right)^{lM}\beta^{l}\right]$$

Now taking the derivative of $Q_0(x,N)$ with respect to $x$ (for this section derivatives are always with respect to $x$) we find:

$$Q_{0'}(x,N)=\frac{(\beta-1)(1-x)^{N-1}}{\beta x^{2}}\sum_{l=1}^{\left\lfloor\frac{N}{M}\right\rfloor}\left[\binom{N-1}{lM-1}\left(\frac{x}{1-x}\right)^{lM}\beta^{l}\big(lM-1-x(N-1)\big)\right] \tag{12}$$

Note then that to determine when $Q_{0'}(x, N) = 0$, we need only examine the sum portion. As such we redefine this sum as $S(x, N)$, where:

$$S(x, N) = \sum_{l=1}^{\left\lfloor \frac{N}{M} \right\rfloor} h_l\,(x, N) \tag{13}$$

$$h_l(x, N) = \binom{N-1}{lM-1}\left(\frac{x}{1-x}\right)^{lM} \beta^l\big(lM - 1 - x(N-1)\big) \tag{14}$$

Notice that each term $h_l(x, N)$ is zeroed by a specific $x$ value, namely:

$$x_l^* = \frac{lM-1}{N-1} \tag{15}$$

And for future reference we will also note,

$$\frac{x_l^*}{1-x_l^*} = \frac{lM-1}{N-lM} \tag{16}$$

Given our interest in finding the zeros of $S(x, N)$, we will use these points $x_l^*$ as possible candidate zeros. We hope to find that for $x_l^* = \frac{lM-1}{N-1}$ then $Q'(x_l^*) \approx 0$. This makes theoretical sense, seeing as this would mean that the change in cooperation rate is highest when the group is one cooperator away from reaching the next threshold, thus creating a strong pressure for wanting to cooperate. We are left with needing to show that this candidate zero is likely to be a zero of equation (6.1.12).

We start by taking a Taylor expansion of $S(x, N)$ centred around $x_l^*$. For simplicity and given that we are particularly interested in the behaviour of $S(x, N)$ around $x_l^*$, we will be working with the first two terms of the Taylor expansion.

$$S(x, N) = S(x_l^*, N) + (x - x_l^*)S'(x_l^*, N) + (x - x_l^*)^2\frac{S''(x_l^*, N)}{2!} + \cdots$$
$$S(x, N) \approx S(x_l^*, N) + (x - x_l^*)S'(x_l^*, N)$$
$$S(x, N) \approx R_l + (x - x_l^*)S'(x_l^*, N)$$

Where,

$$R_l = \sum_{j \neq l} h_j\,(x_l^*, N) \tag{17}$$

And near $x_l^*$, we assume that $h'_l(x, N)$ dominates $S'(x, N)$ and so we make the following substitution,

$$S'(x_l^*, N) \approx h'_l(x_l^*, N)$$

We can check that this holds true, but given the shared structure between the terms of $R_l(x, N)$ and $S'(x, N)$, if we show that $h_{l'}$ dominates $R_l(x, N)$, which we will be demonstrating below, then we will have also shown that $h'_l(x, N)$ dominates $R_l(x, N)$.

Searching for the zeros of this function we get:

$$0 \approx R_l + (x - x_l^*)h'_l(x_l^*, N)$$
$$\delta_l = x - x_l^* \approx -\frac{R_l}{h'_l(x_l^*, N)}$$

Where $\delta_l$ is the distance between our candidate zero $x_l^*$ and a true zero of $S(x_l^*)$. Thus we have that,

$$\delta_l \approx -\frac{R_l}{h'_l(x_l^*)} \approx -\sum_{j \neq l} \frac{h_j(x, N)}{h'_l(x, N)}$$

And we are particularly interested in the absolute distance of the true zero and candidate zero so we will be trying to establish an upper bound on $|\delta_l|$. We start by taking the derivative of $h_l(x, N)$ with respect to $x$.

$$h'_l(x, N) = \binom{N-1}{lM-1}\left(\frac{x}{1-x}\right)^{lM} \beta^l \left(lM\frac{lM - 1 - x(N-1)}{x(1-x)} - (N-1)\right)$$

So, for $h'_l(x_l^*, N)$:

$$h_{l'}(x_l^*, N) = \binom{N-1}{lM-1}\left(\frac{x_l^*}{1-x_l^*}\right)^{lM} \beta^l \left(lM\frac{lM - 1 - (lM-1)}{x_l^*(1-x_l^*)} - (N-1)\right)$$
$$h_{l'}(x_l^*, N) = -\binom{N-1}{lM-1}\left(\frac{lM-1}{N-lM}\right)^{lM} \beta^l (N-1)$$

Next, we find $h_j(x_l^*, N)$.

$$h_j(x_l^*, N) = \binom{N-1}{jM-1}\left(\frac{lM-1}{N-lM}\right)^{jM} \beta^j M(j-l)$$

Thus, we can rewrite each term of our sum as:

$$\frac{h_j(x_l^*, N)}{h'_l(x_l^*, N)} = -\frac{\binom{N-1}{jM-1}\left(\frac{lM-1}{N-lM}\right)^{jM} \beta^j M(j-l)}{\binom{N-1}{lM-1}\left(\frac{lM-1}{N-lM}\right)^{lM} \beta^l (N-1)}$$
$$\frac{h_j(x_l^*, N)}{h'_l(x_l^*, N)} = -\frac{\binom{N-1}{jM-1}}{\binom{N-1}{lM-1}}\left(\frac{lM-1}{N-lM}\right)^{M(j-l)} \beta^{j-l} \frac{M(j-l)}{N-1}$$

And so,

$$\delta_l \approx -\sum_{j \neq l} \frac{h_j(x_l^*, N)}{h'_l(x_l^*, N)} = \sum_{j \neq l} \frac{\binom{N-1}{jM-1}}{\binom{N-1}{lM-1}}\left(\frac{lM-1}{N-lM}\right)^{M(j-l)} \beta^{j-l} \frac{M(j-l)}{N-1}$$

Let us then call the absolute value of the component ratio $F_{j,l}$. Meaning that,

$$|\delta_l| \approx \left|\sum_{j\neq l} \frac{h_j(x_l^*, N)}{h'_l(x_l^*, N)}\right| \leq \sum_{j\neq l} \left|\frac{h_j(x_l^*, N)}{h'_l(x_l^*, N)}\right| = \sum_{j\neq l} F_{j,l}$$

Where,

$$F_{j,l} = \left|\frac{h_j(x_l^*, N)}{h'_l(x_l^*, N)}\right| = \frac{\binom{N-1}{jM-1}}{\binom{N-1}{lM-1}} \left(\frac{lM-1}{N-lM}\right)^{M(j-l)} \beta^{j-l} \frac{M|j-l|}{N-1}$$

We want to find upper bounds on this ratio, first only examining $j$ larger than $l$, or $j \geq l+1$.

$$\frac{F_{j+1,l}}{F_{j,l}} = \left[\frac{\binom{N-1}{(j+1)M-1}}{\binom{N-1}{lM-1}} \left(\frac{lM-1}{N-lM}\right)^{M((j+1)-l)} \beta^{(j+1)-l} \frac{M|(j+1)-l|}{N-1}\right]$$

$$\cdot \left[\frac{\binom{N-1}{lM-1}}{\binom{N-1}{jM-1}} \left(\frac{lM-1}{N-lM}\right)^{-M(j-l)} \beta^{l-j} \frac{N-1}{M|j-l|}\right]$$

$$\frac{F_{j+1,l}}{F_{j,l}} = \frac{\binom{N-1}{(j+1)M-1}}{\binom{N-1}{jM-1}} \beta \left(\frac{lM-1}{N-lM}\right)^{M} \left|\frac{j+1-l}{j-l}\right|$$

To study this, we need to unpack the binomial coefficients. Using $\binom{n}{k} = \frac{n!}{k!(n-k)!}$, we get:

$$\frac{\binom{N-1}{(j+1)M-1}}{\binom{N-1}{jM-1}} = \frac{(N-1)!}{\big((j+1)M-1\big)!\,(N-1-(j+1)M+1)!} \cdot \frac{(jM-1)!\,(N-1-jM+1)!}{(N-1)!}$$

$$\frac{\binom{N-1}{(j+1)M-1}}{\binom{N-1}{jM-1}} = \frac{(jM-1)!}{\big((j+1)M-1\big)!} \cdot \frac{(N-jM)!}{(N-(j+1)M)!}$$

And we notice that a lot of these terms will cancel out, meaning

$$\frac{(jM-1)!}{\big((j+1)M-1\big)!} = \frac{1}{(jM)(jM+1)\cdots\big((j+1)M-1\big)}$$

$$\frac{(N-jM)!}{(N-(j+1)M)!} = (N-jM)(N-jM-1)\cdots(N-(j+1)M+1)$$

Then this can be written as the product of ratios.

$$\frac{\binom{N-1}{(j+1)M-1}}{\binom{N-1}{jM-1}} = \prod_{t=1}^{M} \frac{N-jM-t+1}{jM+t-1}$$

And replacing this above we get,

$$\frac{F_{j+1,l}}{F_{j,l}} = \left[\prod_{t=1}^{M} \frac{N-jM-t+1}{jM+t-1}\right] \beta \left(\frac{lM-1}{N-lM}\right)^{M} \left|\frac{j+1-l}{j-l}\right|$$

$$\frac{F_{j+1,l}}{F_{j,l}} \leq \left(\frac{N-jM}{jM}\right)^M \beta \left(\frac{lM-1}{N-lM}\right)^M \left|\frac{j+1-l}{j-l}\right|$$

As $j$ increases, then the term $\frac{N-jM}{jM}$ decreases, and so the largest this ratio will ever be is at $j = l+1$. Similarly, the largest possible value of $\frac{j+1-l}{j-l}$ is $2$. This means that for all $j \geq l+1$ we can bound $\frac{F_{j+1,l}}{F_{j,l}}$ by $q_+$, where $q_+$ is defined as:

$$\frac{F_{j+1,l}}{F_{j,l}} \leq 2\beta \left(\frac{N-(l+1)M}{(l+1)M}\right)^M \left(\frac{lM-1}{N-lM}\right)^M \coloneqq q_+ \tag{18}$$

Now for $j$ smaller than $l$, or $j \leq l-1$, we will do the same thing to find an upper bound $q_-$, only noting that by symmetry all the ratios are flipped. This means that

$$\frac{F_{j-1,l}}{F_{j,l}} \leq \frac{2}{\beta}\left(\frac{(l-1)M}{N-(l-1)M}\right)^M \left(\frac{N-lM}{lM-1}\right)^M \coloneqq q_- \tag{19}$$

Note that $q_+$ is likely less than $1$. We can see this by rewriting it,

$$q_+ = 2\beta \left(1 - \frac{M}{N-lM}\right)^M \left(\frac{lM-1}{lM+M}\right)^M$$

And $1 - \frac{M}{N-lM} < 1$ and $\frac{lM-1}{lM+M} < 1$. As such $q_+ > 1$ will only occur for large values of $\beta$ or perhaps our largest $l$. Similarly, $q_- < 1$ unless $\beta$ is large or perhaps for our largest $l$. Under the condition where $q_+ < 1$ and $q_- < 1$ are both true, then

$$\sum_{j=l+1}^{\left\lfloor \frac{N}{M} \right\rfloor} F_{j,l} \leq F_{l+1,l} + q_+ F_{l+1,l} + q_+^2 F_{l+1,l} + \ldots \leq \frac{F_{l+1,l}}{1-q_+}$$

And,

$$\sum_{j=1}^{l-1} F_{j,l} \leq \frac{F_{l-1,l}}{1-q_-}$$

Together,

$$|\delta_l| = \sum_{j \neq l} F_{j,l} \leq \frac{F_{l+1,l}}{1-q_+} + \frac{F_{l-1,l}}{1-q_-} := T_l$$

It should also be said that in the future we will require even smaller bounds on $q_+$ and $q_-$, and so cases where these bounds are greater or equal to $1$ fall outside of our analysis, but in such cases we are unlikely to be able to bound $|\delta_l|$. We now have a bound on the absolute distance between our candidate zero and true zero. Next we want to show that there does in fact exist a true zero, no more than $T_l$ away from our candidate zero. Notice that each candidate zero are $\Delta$ apart where,

$$\Delta = x_{l+1}^{*} - x_{l}^{*} = \frac{M}{N-1} \tag{20}$$

Now so long as we have:

$$|\delta_l| < \frac{\Delta}{2} \tag{21}$$

Then we have that each true zero is well spaced pertains to only one candidate zero. That is we would know the true zero will fall within the interval $\left[x_l^* - \frac{\Delta}{2}, x_l^* + \frac{\Delta}{2}\right]$, or in other words there exists a true zero near our candidate zero. In other words, we want to show that our bound on $\delta_l$, $T_l$, is smaller than half the distance between two candidate zeros, Δ, or

$$\frac{F_{l+1,l}}{1-q_+} + \frac{F_{l-1,l}}{1-q_-} < \frac{M}{2(N-1)}$$

Expanding this and replacing the ratio of binomial coefficients with the product found earlier, we get

$$F_{l+1,l} = \frac{\binom{N-1}{(l+1)M-1}}{\binom{N-1}{lM-1}} \left(\frac{lM-1}{N-lM}\right)^{M((l+1)-l)} \beta^{(l+1)-l} \frac{M|(l+1)-l|}{N-1}$$
$$F_{l+1,l} = \left[\prod_{t=1}^{M} \frac{N-lM-t+1}{lM+t-1}\right] \left(\frac{lM-1}{N-lM}\right)^{M} \beta \frac{M}{N-1}$$

Because the smallest numerator possible is $N - lM - M + 1$, for $t = M$ and the largest denominator possible is $lM + M - 1$, for $t = M$. Then we know that the whole product is bounded below by $\left(\frac{N-lM-M+1}{lM+M-1}\right)^M$, which is itself larger than $\left(\frac{N-(l+1)M}{(l+1)M}\right)^M$, then we have

$$F_{l+1,l} \geq \left(\frac{N-(l+1)M}{(l+1)M}\right)^{M} \left(\frac{lM-1}{N-lM}\right)^{M} \beta \frac{M}{N-1}$$
$$F_{l+1,l} \geq \frac{1}{2} q_+ \left(\frac{M}{N+1}\right)$$

And similarly,

$$F_{l-1,l} = \frac{\binom{N-1}{(l-1)M-1}}{\binom{N-1}{lM-1}} \left(\frac{lM-1}{N-lM}\right)^{M((l-1)-l)} \beta^{(l-1)-l} \frac{M|(l-1)-l|}{N-1}$$
$$F_{l-1,l} = \left[\prod_{t=1}^{M} \frac{lM-t}{N-lM+t}\right] \left(\frac{lM-1}{N-lM}\right)^{-M} \frac{1}{\beta} \frac{M}{N-1}$$
$$F_{l-1,l} \geq \left(\frac{(l-1)M}{N-(l-1)M}\right)^{M} \left(\frac{lM-1}{N-lM}\right)^{-M} \frac{1}{\beta} \frac{M}{N-1}$$

$$F_{l-1,l} \geq \frac{1}{2} q_- \frac{M}{N-1}$$

So together,

$$\frac{1}{2}q_+\frac{M}{N-1}\left(\frac{1}{1-q_+}\right)+\frac{1}{2}q_-\frac{M}{N-1}\left(\frac{1}{1-q_-}\right)\le\frac{F_{l+1,l}}{1-q_+}+\frac{F_{l-1,l}}{1-q_-}<\frac{M}{2(N-1)}$$

Or,

$$\frac{q_+}{1-q_+}+\frac{q_-}{1-q_-}\le 1 \tag{22}$$

Next, we will further simplify these terms.

$$q_+ = 2\beta\left(\frac{N-(l+1)M}{(l+1)M}\right)^M\left(\frac{lM-1}{N-lM}\right)^M$$
$$q_+ = 2\beta\left(\frac{(N-lM-M)(lM-1)}{(lM+M)(N-lM)}\right)^M$$
$$q_+ = 2\beta A_l^M \tag{23}$$

Where,

$$A_l=\frac{(N-lM-M)(lM-1)}{(lM+M)(N-lM)} \tag{24}$$

And,

$$q_- = \frac{2}{\beta}\left(\frac{(l-1)M}{N-(l-1)M}\right)^M\left(\frac{lM-1}{N-lM}\right)^{-M}$$
$$q_- = \frac{2}{\beta}\left(\frac{(lM-M)(N-lM)}{(N-lM+M)(lM-1)}\right)^M$$
$$q_- = \frac{2}{\beta}B_l^M \tag{25}$$

Where,

$$B_l=\frac{(lM-M)(N-lM)}{(N-lM+M)(lM-1)} \tag{26}$$

And,

$$A_lB_l=\frac{(N-lM-M)(lM-M)}{(lM+M)(N-lM+M)}$$
$$A_lB_l=\frac{(u-M)(v-M)}{(v+M)(u+M)}\coloneqq C_l \tag{27}$$

Where $u=N-lM$ and $v=lM$. Meaning,

$$q_+q_- = 4C_l^M \tag{28}$$

And so long, as $u>M$ and $v>M$ (or $1<l<\frac{N}{M}$), then

$$0<C_l=\frac{(u-M)(v-M)}{(v+M)(u+M)}<1$$

These bounds on $u$ also give us that $\frac{u-M}{u+M} < 1$ and so,

$$q_+ q_- = 4C_l^M = 4\left(\frac{u-M}{u+M}\right)^M \left(\frac{v-M}{v+M}\right)^M < 4\left(\frac{v-M}{v+M}\right)^M = 4\left(\frac{l-1}{l+1}\right)^M$$

Next we further bound $q_+$ and $q_-$, so

$$A_l = \frac{(N-lM-M)(lM-1)}{(lM+M)(N-lM)}$$
$$A_l < \frac{(N-lM)(lM)}{(lM+M)(N-lM)}$$
$$A_l < \frac{l}{l+1}$$
$$q_+ < 2\beta\left(\frac{l}{l+1}\right)^M \quad (29)$$

Similarly,

$$B_l = \frac{(lM-M)(N-lM)}{(N-lM+M)(lM-1)}$$
$$B_l < \frac{(lM-M)(N-lM)}{(N-lM)(lM-1)}$$
$$B_l < \frac{lM-M}{lM-1}$$
$$B_l < \frac{l-1}{l-\frac{1}{M}}$$
$$q_- < \frac{2}{\beta}\left(\frac{l-1}{l-\frac{1}{M}}\right)^M \quad (30)$$

Finally recalling that we actually want to study the inequality (22)

$$\frac{q_+}{1-q_+} + \frac{q_-}{1-q_-} < 1$$

This is can be true for a number of combinations of values for $q_+$ and $q_-$, but will always be true if both $q_+ < \frac{1}{3}$ and $q_- < \frac{1}{3}$. So combining all of our bounds, if we show that the following 3 conditions hold,

$$2\beta\left(\frac{l}{l+1}\right)^M < \frac{1}{3}; \qquad \frac{2}{\beta}\left(\frac{l-1}{l-\frac{1}{M}}\right)^M < \frac{1}{3}; \qquad 4\left(\frac{l-1}{l+1}\right)^M < \frac{1}{9}$$

Then we will have shown that

$$|\delta_l| \le T_l < \frac{\Delta}{2}$$

Meaning that there will exist a true zero centered around each of our candidate zeros $x_l^*$. What is left is to check when these three conditions hold. Starting with edge cases $l = 1$ and $l = \left\lfloor \frac{N}{M} \right\rfloor$, noting that in this case the assumptions needed to make $q_+ q_- = 4C_l^M$ do not hold and so it does not need to be checked. For $l = 1$, $q_- = 0$ so we only need to check $q_+$.

$$
\begin{aligned}
q_+ \leq 2\beta\left(\frac{1}{2}\right)^M &< \frac{1}{3} \\
\left(\frac{1}{2}\right)^M &< \frac{1}{6\beta} \\
M &> \frac{\ln(6\beta)}{\ln(2)}
\end{aligned}
\tag{31}
$$

In the main manuscript we typically set $\beta = 2$ which under these conditions would represent $M > \frac{\ln 12}{\ln 2} \approx 3.6$. Meaning so long as $M \geq 4$, then there should be a zero near $x_1^*$. Larger multipliers increase the required threshold. If $\beta = 3$, then $M > \frac{\ln 18}{\ln 2} \approx 4.2$ and so $M \geq 5$ is required to have a zero near $x_1^*$.

The other edge case is $l = \left\lfloor \frac{N}{M} \right\rfloor$, then $q_+ = 0$ and,

$$
\begin{aligned}
q_- \leq \frac{2}{\beta}\left(\frac{\frac{N}{M} - 1}{\frac{N}{M} - \frac{1}{M}}\right)^M &< \frac{1}{3} \\
\frac{2}{\beta}\left(\frac{N - M}{N - 1}\right)^M &< \frac{1}{3} \\
\left(\frac{N - M}{N - 1}\right)^M &< \frac{\beta}{6}
\end{aligned}
\tag{32}
$$

This boundary is less straight forward but notice that as $M$ increases then the LHS decreases, making the inequality more likely to hold. On the flip side as $N$ increases then the LHS increases and the inequality is less likely to hold. To analyse this inequality we will look at the ratio between $N$ and $M$ required to make the inequality hold, where we set $N = rM$, for some $r > 1$.

$$
\begin{aligned}
\left(\frac{rM - M}{rM - 1}\right)^M &< \frac{\beta}{6} \\
\left(\frac{M(r-1)}{rM}\right)^M < \left(\frac{M(r-1)}{rM - 1}\right)^M &< \frac{\beta}{6} \\
\left(\frac{r-1}{r}\right)^M &< \frac{\beta}{6} \\
r &< \frac{1}{1 - \left(\frac{\beta}{6}\right)^{\frac{1}{M}}}
\end{aligned}
\tag{33}
$$

So, we have a bound on the ratio between $N$ and $M$. To understand what this boundary means, we will set some parameter values and examine the result, using $\beta = 2$ as before. If $M = 4$ (our earlier bound), then $N \leq 16$, but an even slightly higher $M = 6$, then $N \leq 35$ and the much higher $M = 12$, then $N \leq 137$. So, when the thresholds are very closely spaced (small $M$), then this inequality is unlikely to hold. Whereas further spaced thresholds are stable for much larger populations. Increasing the multiplier $\beta$ increases the tenable population size, for instance with $\beta = 3$, then a threshold of $M = 6$ requires $N \leq 54$. This means that larger thresholds can support larger populations while ensuring our inequalities hold.

Finally, we have the general case for $l$, where both bounds are relevant. For these we will study them using $4\left(\frac{l-1}{l+1}\right)^M < \frac{1}{9}$. As it does not require $\beta$ to evaluate.

$$
\begin{aligned}
4\left(\frac{l-1}{l+1}\right)^M &< \frac{1}{9} \\
\left(\frac{l-1}{l+1}\right)^M &< \frac{1}{36} \\
\frac{l-1}{l+1} &< 36^{\frac{-1}{M}} \\
l &< \frac{1+36^{\frac{-1}{M}}}{1-36^{\frac{-1}{M}}}
\end{aligned} \tag{34}
$$

Here as $M$ increases then $l$ values which fall below the bound also increase. Recalling that in this case we are studying $2 \le l \le \left\lfloor \frac{N}{M} \right\rfloor - 1$. So for a given $M$ value, this will place an upper bound on possible $l$ values for which the inequalities hold, which in turn places an upper bound on $N$ values for which the inequality holds. Some examples include $M = 4$, for which $l \le 2$ holds and $N \le 13$ holds. $M = 6$, for which $l \le 3$ holds and $N \le 26$ holds, and $M = 12$, for which $l \le 6$ holds and $N \le 92$ holds.

Taking these pieces of information together, we find that the general trend is that so long as the thresholds are sufficiently spaced apart then there will exist a local maxima in cooperation change at each threshold, right before the threshold is met ($x_l^* = \frac{lM-1}{N-1}$). What counts as sufficiently spaced depends on the size of the population. In a group of $N = 20$ then a threshold at $M = 10$ is quite daunting, but in a group of $N = 200$ a threshold of $M = 10$ is quite small and attainable with only a slight change in cooperation rate. As such, cooperation change will keep increasing and have no local maxima at $x_l^*$. We can use our bounds listed above as a baseline check for knowing whether there is likely to be a maxima for a set of parameters, recalling that these bounds are likely conservative estimates of the true upper bounds.

Altogether this means we have that $|x - x_l^*| \approx \frac{|S(x_l^*,N)|}{|S'(x_l^*,N)|} \approx 0$. Meaning that $x_l^*$ is an approximate zero of $S(x,N)$, which in turn means that $x_l^*$ is an approximate zero of our larger function $Q'_0(x,N)$, and an approximate extremum of $Q_0(x,N)$.

Finally, we need to take the second derivative of $Q_0(x,N)$, with respect to $x$ evaluated at $x = x_l^*$ to check that $x_l^*$ is in fact a local maximum. First notice the following,

$$
\begin{aligned}
Q_{0'}(x,N) &= \frac{(\beta-1)(1-x)^{N-1}}{\beta x^2} S(x,N) \\
Q_{0''}(x,N) &= -\frac{(\beta-1)(1-x)^{N-2}}{\beta x^3}\Big(S(x,N)\big((N-3)x+2\big) - x(1-x)S'(x,N)\Big)
\end{aligned}
$$

Recalling that at $S(x_l^*,N) \approx h_l(x_l^*,N) = 0$ and $S'(x_l^*,N) \approx h'_l(x_l^*,N)$.

$$
\begin{aligned}
Q_{0''}(x_l^*,N) &= -\frac{(\beta-1)(1-x_l^*)^{N-2}}{\beta {x_l^*}^3}\Big(S(x_l^*,N)\big((N-3)x_l^*+2\big) - x_l^*(1-x_l^*)S'(x_l^*,N)\Big) \\
Q_{0''}(x_l^*,N) &\approx \frac{(\beta-1)(1-x_l^*)^{N-2}}{\beta {x_l^*}^3}\big(x_l^*(1-x_l^*)h'(x_l^*,N)\big)
\end{aligned}
$$

And since $0 < x_l^* < 1$ and $\beta > 1$, the sign of $Q_{0''}(x_l^*, N)$ is determined by the sign of $h'(x_l^*, N)$ and earlier we found that

$$h_{l'}(x_l^*, N) = -\binom{N-1}{lM-1}\left(\frac{lM-1}{N-lM}\right)^{lM} \beta^l (N-1) < 0$$

Which implies $Q''_0(x_l^*, N) < 0$, meaning that $x_l^*$ is approximately a near maximum of $Q_0(x, N)$. Altogether this means that cooperation change has a local maximum at approximately $x_l^* = \frac{lM-1}{N-1}$, or when the group is likely to be one cooperator away from a next threshold, so long as the thresholds are sufficiently distanced. This local maxima may not exist for large $\beta$ values or small $M$ values. In both of these cases the drive for increasing cooperation remains too strong even after a threshold is met so cooperation will continue to increase. We see this behaviour in Figure S1**Error! Reference source not found.**, including the edge case where $M$ is too small for this pattern to hold.

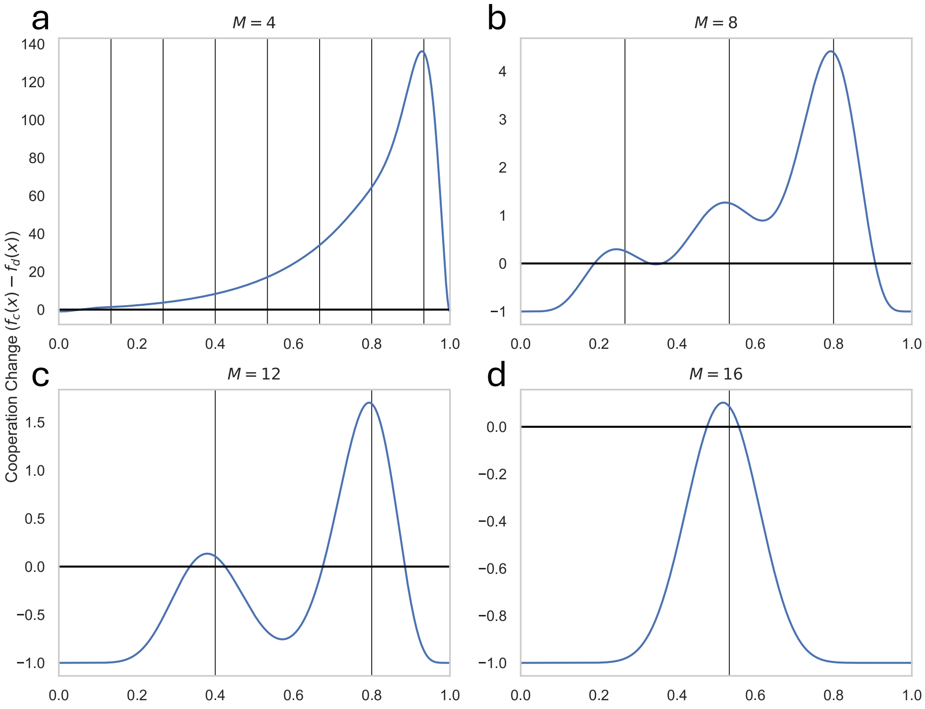


**Figure S1: Cooperation Change For Multiple Thresholds**
Figure showing the change in cooperation rate for different $\boldsymbol{M}$ values. Notice that the plots for $\boldsymbol{M = 8}$ and $\boldsymbol{M = 12}$ (**(b)** and **(c)**) appear in Figure 1 of the manuscript, with $\boldsymbol{x_l^*}$ highlighted by vertical lines. When $\boldsymbol{M}$ is too small, as per our conditions listed above, then $\boldsymbol{x_l^*}$ stops being a local maxima and instead cooperation change continues to rise. This is the case for $\boldsymbol{M = 4}$ **(a)**. (Parameters: group size $\boldsymbol{N = 30}$, cooperation cost $\boldsymbol{c = 1}$, cooperation benefit $\boldsymbol{F = 225}$, threshold multiplier $\boldsymbol{\beta = 2}$)

We do not show where the local minima of the function occur, but by the Intermediate Value Theorem there must be at least one local minimum in between each $x_l^*$, or in between thresholds.

This analysis has only been done when there is no effect of technology in our model. Otherwise, our rewriting of $Q(x, N)$ would not work. We can speculate that including technology would push the peaks of our function to the left, essentially making cooperation increases occur earlier since each threshold is attainable with fewer people. I suspect that thresholds will occur roughly at $x_l^* = \frac{lM-1}{N^{1+\lambda}-1}$, which also holds the pattern of our previous analysis for $\lambda = 0$, but I will not show that here.

**Analyzing Population Change**

Doing the same sort of analysis as we did for cooperation change, but this time for population change. Here we note that population growth is dictated by the term,

$$g\big(U(x,N)\big) - N \tag{35}$$

For simplicity we will once more set $\lambda = 0$. And we want to know when its derivative with respect to $N$ is zero, or

$$1 = g'\big(U(x,N)\big)$$
$$1 = \alpha U'(x,N)$$
$$1 = \alpha\left(x\frac{d}{dN}f_C(x,N) + (1-x)\frac{d}{dN}f_D(x,N)\right)$$

Due to $N$ denoting population size, we can only take an approximate discrete derivative of $f_C$ and $f_D$ with respect to $N$. As such, we approximate this derivative as the following:

$$\frac{d}{dN}f_C(x,N) \approx f_C(x,N+1) - f_C(x,N) \tag{36}$$
$$\frac{d}{dN}f_D(x,N) \approx f_D(x,N+1) - f_D(x,N) \tag{37}$$

We could try and further simplify these equations, however we are unable to find a similar bound on parameter values like we did for cooperation rate change and so it is unlikely to be beneficial. Instead, the best we can do is verbally describe these functions and try to extract a pattern. The key question becomes, if we add one extra person to the group, does that increase or decrease the group utility.

If the group is very close to a threshold, note that here this is when $xN \approx lM$ because we will have roughly $xN$ cooperators in the group, then adding one person is likely to increase the utility because if they cooperate then utility will greatly increase. This means that the effect of adding a person to the utility is greater when the cooperation rate is high. Adding an extra defector is unlikely to do anything. The benefit of adding additional people also increases over time, because each new threshold is more valuable than the last. As such we expect the local maxima of population change to occur slightly past a threshold, perhaps closer to $xN \approx lM + c$, but for $c$ which decreases with each threshold. If the group is far from a threshold, then the utility decreases by adding extra people because this lowers everyone's utility.

Taking this with the cooperation change work, we have that the rate of cooperation change peaks right before a threshold is met and the rate of population change peaks right after a threshold is met. The fact that these peaks occur on either side of the threshold is what makes the ratcheting effect of our model effective. Being near a threshold means an increase in cooperation rate which means going past a threshold which means an increase in population, and in turn this can mean being once more near a threshold.

**Both Variables**

Allowing both cooperation rate and population to vary, we can also find when these are both stable. That is to say, when is

$$0 = x(1-x)\big(f_C(x,N) - f_D(x,N)\big)$$
$$0 = rN\frac{g\big(U(x,N)\big) - N}{g\big(U(x,N)\big)}$$

If we only take the interior zeros of cooperation change, we are left with,

$$f_C(x,N) = f_D(x,N) \tag{38}$$
$$N = g\big(U(x,N)\big) \tag{39}$$

And making the substitution from equation (6.1.38) into equation (6.1.39) we are left with:

$$N = N_0 + \alpha\big(xf_C(x,N) + (1-x)f_D(x,N)\big)$$
$$N = N_0 + \alpha f_D(x,N)$$
$$f_D(x,N) = \frac{N - N_0}{\alpha} \tag{40}$$

Again, we will not fully analyze but we will describe some of the behavior. We do so by varying $N$ and solving for $x$ such that equation (40) is satisfied. Here we generally find that cooperation peaks for small populations and decreases as population grows. Mathematically this occurs because increasing $N$ grows the $RHS$ linearly, but the $LHS$ grows exponentially through greater cooperative returns. As such, to balance the growth on the $LHS$ a smaller cooperation rate ($x$) is necessary, or a larger decay in returns with an increasing population through our decay parameter ($\rho$). In terms of our model, when the population increases then some new cooperators are added to the group, and so the benefits of cooperation can be maintained with a slightly lower cooperation rate. For instance, if $N = 10$, then having $8$ cooperators corresponds to $x = 0.8$, but when $N = 11$, then $8$ cooperators corresponds to $x = 0.73$. For this same reason, introducing resource constraints increases the cooperation rate. For one it increases the $RHS$ and so the $LHS$ can be balanced by slightly increasing the cooperation rate. In practice this means that because having an increase in population reduces the effect of average utility on population growth, this must be compensated for by increasing cooperation.

Technology effects the model by decreasing cooperation required for a stable equilibrium. This may seem counter intuitive, but it allows cooperation to be stable with fewer cooperators. Essentially, this greatly increases the $LHS$ and so the $LHS$ needs to be decreased by having a lower cooperation rate to keep the equation equal. This means the stable cooperation rate to stabilize population will be lower. If we do not force both cooperation rate and population size to be stable, then we should instead expect technology to lead to a big increase in cooperation because further thresholds will be more attainable. This would lead to a constant increase in population and cooperation.

Resource decay effects the model by increasing the cooperation required for a stable equilibrium. It adds a pressure to the group to reach thresholds with fewer people, thus leading to smaller but more cooperative groups. It does however make it more difficult to hit subsequent thresholds, thus perhaps preventing the scaling of cooperation.

**Linear Growth**

If we consider non-super linear growth in cooperative returns, then the same spikes and troughs behavior exists, but the maximum value of different spikes will be roughly equal. This is evident in Figure S2 where the difference between any two thresholds is fixed and so cooperation change doesn't increase across thresholds. As such, although the general pattern of peaks and troughs persist, the ratcheting mechanic which allows for population growth breaks down. This tells us that for cooperation to expand there must be super linear growth. Similarly, the expansion occurs quicker the larger the growth term.

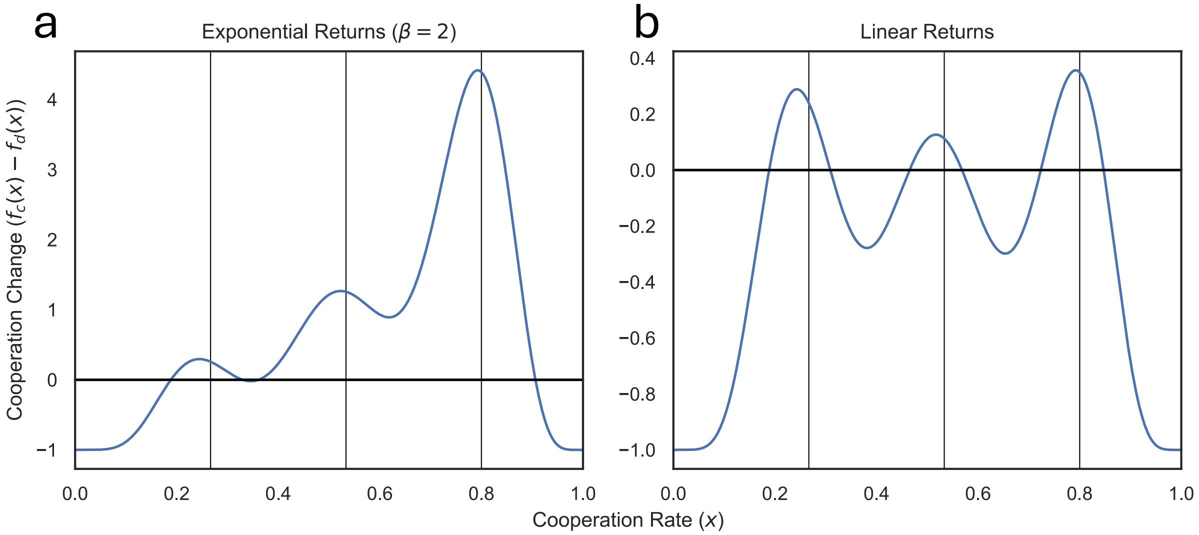


**Figure S2: Linear versus Super Linear Growth**
Cooperation change given super linear returns **(a)** versus linear returns **(b)**. Thresholds still shape cooperative behavior in the linear case, but the difference between thresholds is not as pronounced. This is due to diminishing relative returns of cooperation. Regardless of threshold reached, the increase in cooperative returns from one threshold to the next is always $\frac{\boldsymbol{Fc}}{\boldsymbol{N}}$. As such, there is no material difference in returns from going from the first to the second threshold or the second to third threshold. As such, although there is still reason to reach new thresholds, we no longer see the ratcheting effect and subsequent cooperative growth, as in the super linear condition. (Parameters: group size $\boldsymbol{N} = \mathbf{30}$, threshold size $\boldsymbol{M} = \mathbf{8}$, cooperation cost $\boldsymbol{c} = \mathbf{1}$, cooperation benefit $\boldsymbol{F} = \mathbf{225}$)

A similar case to this, which we do not deeply analyze, is one where the distance between thresholds also grows super linearly. We do this by modifying the cooperative portion of the returns to be:

$$\beta^{\lfloor \log_M(kN^\lambda) \rfloor} - 1 \tag{41}$$

Or $0$ if $k = 0$. Here we find the same sort of effect, where the growth of returns must increase at a faster rate than the growth of distance between thresholds for the cooperative growth effect to occur. That is, the ratcheting effect only occurs if $\beta > M$. We can see some examples of this in Figure S3.

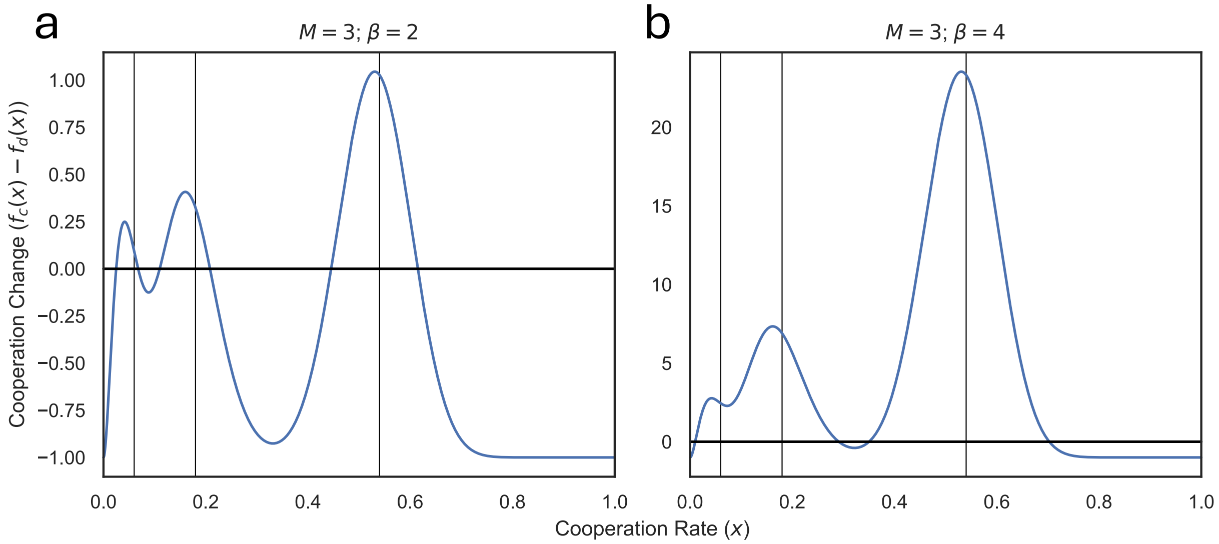


**Figure S3: Non-Linearly Separated Cooperation Thresholds**

Cooperation change graph for non-linearly spaced thresholds, as described in equation (41). In **(a)**, the distance between thresholds ($\boldsymbol{M}$) is greater than the increase in reward between thresholds ($\boldsymbol{\beta}$). In **(b)**, the distance between thresholds is less than the increase in reward between thresholds. As such, in **(a)** we find that cooperation growth appears to stall as the benefit from reaching new thresholds is not as pronounced as the amount of cooperative effort it takes to get there. In **(b)**, the same cooperative growth pattern we found in the original model is roughly replicated, but with thresholds more distantly spaced. As such, we expect that the same ratcheting effect should be possible, although likely requiring larger population growth. (Parameters: group size $\boldsymbol{N = 50}$, cooperation cost $\boldsymbol{c = 1}$, cooperation benefit $\boldsymbol{F = 225}$)

Further analysis of this condition could be worthwhile, as it is unrealistic to say that the difference in cooperation needed to transition from a hunting lifestyle to an agricultural one is the same in terms of manpower as the transition from coal mining to oil drilling. However, for the purposes of the current work, both models appear to behave in a similar manner, and linearly spaced thresholds are much cleaner to interpret and study.
